# Bioprinted Human Primary Arteries Recapitulate Inflammatory Activation and Pharmacologic Rescue

**DOI:** 10.64898/2026.08.14.744906

**Authors:** Zhouquan Fu, V. Amanda Fastiggi, Angelica Phelan, Kailey Bell, Serena Lucarelli, Sarah S. Wilson, John M. Lindner, Alicia A. Cutler

## Abstract

Chronic inflammation drives persistent systemic cytokine signaling that contributes to vascular dysfunction and secondary vasculitis, yet mechanistic studies are limited by models that fail to capture the multicellular architecture and dynamics of human arteries. In contrast, perfusing intact vessels *ex vivo* has limited tractability because of material availability and difficulty of genetic or biochemical manipulation. We developed a modular, perfused artery-on-a-chip platform by tri-axially bioprinting primary human vascular cells to recapitulate the concentric organization of the intimal, medial, and adventitial layers. The engineered vessels are viable longer than 21 days, with functional endothelial barriers, contractile smooth muscle behavior, and actively remodeled extracellular matrices bearing hallmarks of native vascular tissue. Addition of tumor necrosis factor alpha (TNFα) induces altered transcript levels of proinflammatory mediators and secretion of cytokines and matrix-remodeling enzymes without compromising vessel viability. Importantly, this secretory response is effectively attenuated by both a small-molecule JAK1 inhibitor (ABT-317) and anti-TNF antibody (Infliximab), demonstrating the model’s utility for therapeutic evaluation.

## INTRODUCTION

Inflammatory signals in tissue drive the release of proinflammatory cytokines which spread throughout the body via the vasculature. Under chronic inflammatory conditions, these signals are persistently elevated, leading to widespread endocrine responses across many organs and tissues. The vasculature is particularly susceptible, as it both responds to and further amplifies these inflammatory signals, driving endothelial dysfunction and potentially progression to atherosclerosis or broader cardiovascular disease.

Vascular inflammation therefore has a high coincidence with chronic inflammatory conditions. Coincidence of vasculitis with autoimmune diseases ranges from 11 to 56%^1–4^, with evidence that autoimmune pathology precedes vasculitis onset^4^. Similarly, individuals with diabetes mellitus or periodontal disease are at risk of developing vascular inflammation and subsequent atherosclerosis with an increased chance of serious cardiovascular complications^5–8^. Together, these observations emphasize that persistent systemic inflammatory signaling contributes to secondary vasculitis which can affect macro- and microvasculature across the body^10^. In this context, vascular inflammation can develop within a complex environment shaped by interactions among cell types, baseline inflammation, and therapeutic interventions.

Cells in each arterial layer respond to inflammatory signaling and secrete cytokines and chemokines that promote immune cell infiltration^11,12^, further amplifying the initial signal and driving both tissue injury and vascular remodeling. In the intimal layer, pro-inflammatory cytokines disrupt endothelial barrier integrity and induce adhesion molecule expression promoting leukocyte recruitment, tethering, and transmigration into the vessel wall^13,14^. In the medial layer, SMCs shift away from a contractile phenotype and toward a synthetic, proliferative state that results in hyperplasia and vessel stenosis and drives extracellular matrix (ECM) remodeling that weakens the vessel wall^17–19^. In the adventitial layer, fibroblasts also contribute to structural remodeling through ECM deposition and matrix metalloprotease (MMP)– mediated degradation as they transition toward myofibroblast phenotype and shift expression profile toward fibrosis^20–23^. Together, these inflammation-driven changes lead to endothelial dysfunction, vessel wall thickening and fibrosis, lumenal stenosis, and reduced vascular function. Elucidating these dynamics requires model systems that capture the vessel as an interconnected, evolving microenvironment.

Microfluidic vessel-on-a-chip platforms have emerged as powerful *in vitro* systems to recapitulate key aspects of vascular physiology that are poorly captured by conventional two-dimensional cultures or difficult to access in animal models. Ongoing efforts to generate representative 3D vascular models have largely focused on microvasculature using endothelialized microchannels embedded in hydrogels, enabling more mechanistic studies of barrier function, angiogenesis, microvascular transport, and endothelial inflammatory signaling primarily under steady-state flow conditions^1–6^. While instrumental for microvascular biology, these models typically lack the multilayered structure, cellular heterogeneity, and three-dimensional (3D) mechanical environment of arteries^7^. More recent artery-on-a-chip platforms have sought to address these limitations by incorporating additional vascular cell types but largely continue to rely on cell lines and synthetic membranes to segregate co-cultured cells.

To address the need for a more organotypic, scalable complex arterial model, we developed a modular bioprinted vessel and chamber device with the characteristic concentric architecture of native arteries using artery-derived human primary cells. The vessel is maintained under pump-driven pulsatile perfusion, intentionally designed for both continuous monitoring and endpoint collection, and scalable to accommodate the variability of primary human cells. Perturbing this system, we observed a time-dependent response to inflammatory stimulus that was attenuated by treatment with either a small molecule JAK inhibitor or anti-TNF antibody. These features establish this model as a versatile and physiologically relevant platform to study vascular inflammation and therapeutic intervention.

## RESULTS

### Vessel generation and process optimization

We established a sequential biofabrication workflow capable of printing perfusable, tri-layered vessels that accurately model the spatial and cellular distribution of native arteries (**Figure1a).** This process includes initial primary cell selection and bioink preparation as well as bioprinting parameter and post-print processing optimization to balance mechanical integrity, cell morphology, and long-term model viability.

To recapitulate the multi-layered architecture of the native artery with high geometric precision and mechanical stability, we established a multi-step biofabrication strategy utilizing a customized tri-axial bioprinting approach (**Video S1**). The primary cells were encapsulated in a composite bioink consisting of 2% (w/v) alginate, 3% (w/v) gelatin, and 0.5 mg/mL collagen type I at a cell density of 1×10^7^ cells/mL. This specific formulation was engineered to balance printability with a permissive microenvironment for cellular remodeling. The bioinks were loaded into a tri-axial nozzle according to their physiological spatial arrangement, with 30% (w/v) Pluronic F-127 with 100mM CaCl_2_ serving as a sacrificial core to define the lumenal space, the SMC-laden bioink in the middle nozzle, and fibroblast-laden bioink in the outermost nozzle. The printed vessels were extruded using an Allevi 3 bioprinter under optimized pneumatic pressure and deposited directly into an initial crosslinking bath containing CaCl_2_ and transglutaminase (TG). This dual-crosslinking approach provided instantaneous ionic crosslinking of the alginate network to preserve high-fidelity tubular geometry. Following passive removal of the sacrificial Pluronic core during the initial crosslinking step, the vessels underwent a secondary, crosslinking phase with TG to covalently crosslink the gelatin scaffold. For functional maturation, the bioprinted arteries were cannulated within a custom perfusion housing and cannulation sites secured with agarose. To increase the porosity and facilitate cellular interaction with the extracellular matrix (ECM), the alginate component was selectively leached using calcium-free DPBS for 20 minutes. After 48 hours of culture, ECs were seeded into the lumen and statically incubated for an additional 24 hours to allow for EC attachment before integrating the chips into a continuous perfusion system for long-term biomimetic culture. Cells remained viable throughout the extended fabrication process (**Figure 1b**).

**Figure 1:**
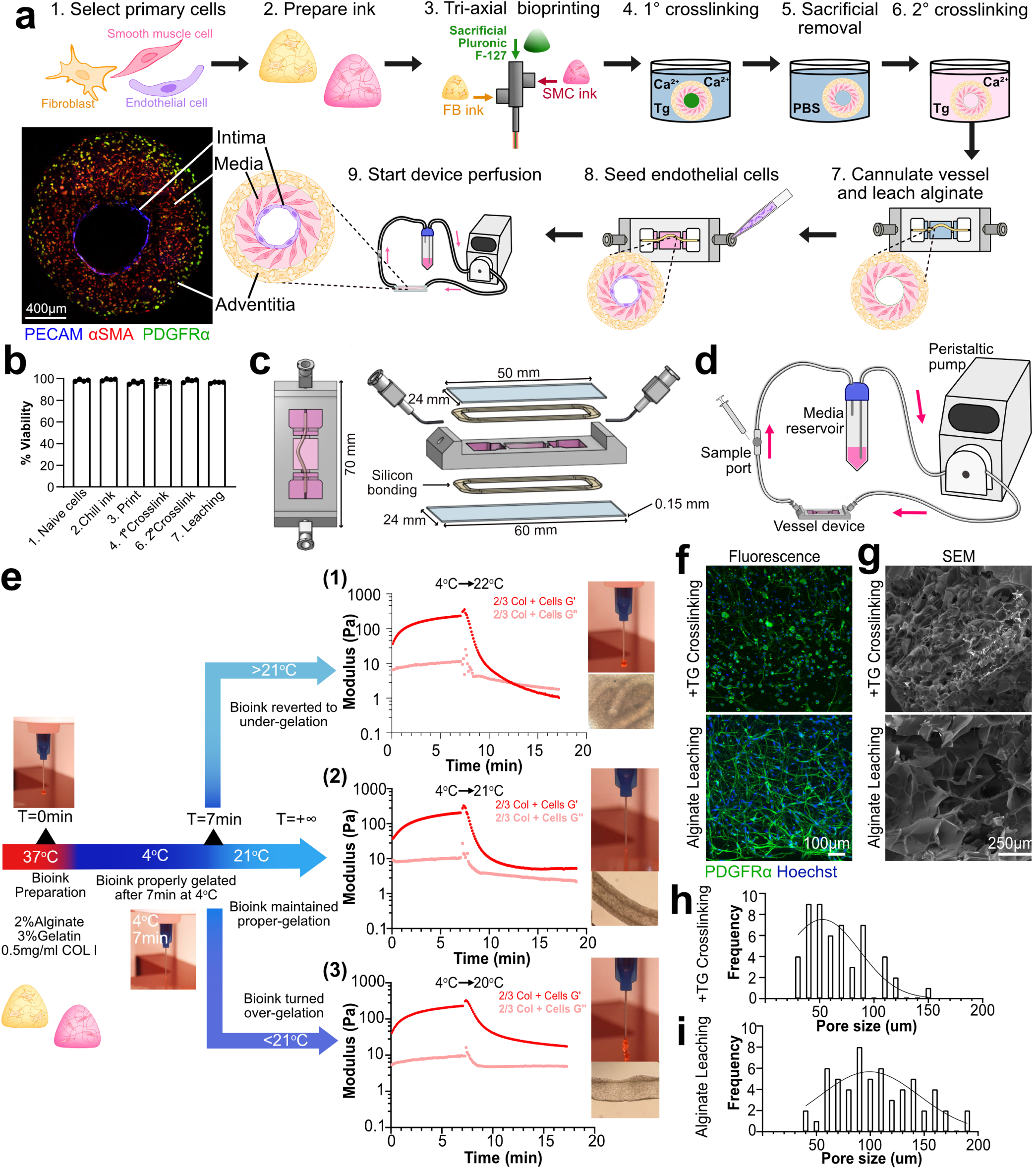
Biofabrication of a three-layer vessel. **(a)** Biofabrication schematic of artery-on-a-chip, including primary cell selection, bioink preparation, tri-axial bioprinting, crosslinking, alginate leaching, and artery assembly on chip, **(b)** Cellular viability throughout the biofabrication workflow quantified by Live (Calcein AM)/Dead (ethidium homodimer-1) dye (mean ± SD, one-way ANOVA, n=4). **(c)** Top and exploded view of the artery-on-a-chip device including chip body, fast curing silicone sealants, bottom and top coverglass, and insert ports, **(d)** Overview of perfusion system with multi­channel peristaltic pump, sampling port, media reservoir, and chip device, **(e)** Schematic of rheology guided tri-axial bioprinting strategy by tunning bioink incubation temperature and time, **(f)** Representative fluorescence microscopy and **(g)** scanning electron microscopy (SEM) images of cell laden hydrogels with and without alginate leaching. Pore size distribution in hydrogels before **(h)** and after **(i)** alginate leaching.

We engineered custom perfusion housing devices designed for compatibility with live imaging, long-term culture, and real-time monitoring along with simple terminal sample recovery (**Figure 1c**). The devices feature external dimensions of a standard microscope slide and top and bottom glass coverslips affixed with rapid-curing silicone adhesive to ensure broad imaging compatibility. The devices maintained structural integrity without leakage for up to 21-days of culture under continuous peristaltic flow. The inside of the device was subdivided into cannulation and working zones. The cannulation sites were secured using 10% (w/v) agarose, providing a high-strength seal for the artery around the cannulae. The central chamber containing the printed vessel was embedded in 5% (w/v) agarose, which provided necessary structural support while maintaining a permissive environment for nutrient exchange. The fluidic circuit was configured as a closed-loop system, featuring an independent media reservoir and dedicated sampling port for sterile cytokine/drug injection and withdrawal of perfusate during longitudinal studies (**Figure 1d**).

FBs and SMCs dynamically transition through cell states that can appear similar and are sensitive to culture conditions. This sensitivity is especially apparent in primary cells. To ensure that we incorporated high quality cells into our prints, we screened commercially purchased primary human arterial FBs, SMCs, and ECs via immunocytochemistry for cell type markers in 2D (**Figure S1a)** and 3D culture (**Figure S1b-c**). While the FB lots expressed both αSMA and PDGFRα in 2D culture, most but not all lots (3 out of 5) decreased αSMA expression after seven days in 3D culture. Furthermore, SMC lots (5 out of 6) maintained αSMA and MYH11 when switched from 2D to 3D culture suggesting that our bioink composition supports the maintenance of healthy FB and SMC phenotypes. To identify optimal FB and SMC lots for printing, we performed RNA sequencing (RNA-seq) of the pre-screened cell lots after 7 days in 3D culture (n = 3 FB donors; n = 5 SMC donors). The cell lots were then organized by hierarchical clustering. We observed notable lot-to-lot heterogeneity, including two cell lots that clustered with the other cell type (**Figure S1d).** We proceeded with cell lots that passed both characterization steps. These findings reinforce the phenotypic plasticity of FBs and SMCs and emphasize the need for thorough molecular characterization before incorporation into complex models.

To evaluate the printability and structural fidelity of the bioink during extrusion-based tri-axial printing, we performed comprehensive rheological characterization of the composite ink both with and without cell encapsulation. Temperature sweep analysis identified a characteristic solid-gel transition at approximately 21°C for the acellular ink (**Figure S2a**) and 18°C for the cell laden ink, indicating that cells within the hydrogel matrix hinder the kinetics of the gelatin random coil-to-triple helix transition. Both formulations exhibited pronounced shear-thinning properties, essential for minimizing cell-damaging shear stress during extrusion where the apparent viscosity decreased by 2-3 orders of magnitude with increasing shear rates from 0.1 s^−1^ to 100 s^−1^ (**Figure S2b**). Notably, the apparent viscosity was lower in the cell-laden bioink compared to its acellular counterpart, potentially due to the displacement of the polymer network by the cellular fraction or altered chain entanglements. Furthermore, as the physical crosslinking of gelatin is a kinetic, time-dependent process, we evaluated the evolution of the storage modulus (G’) at 4°C and 21°C. At 4°C, the G’ exhibited a rapid increase, reaching a stable plateau within 15 minutes, while at 21°C, the storage modulus failed to reach a plateau even after 30 minutes of incubation (**Figure S2c**). For the stability and consistency of our printed vessels, we integrated temperature and time control directly into the printing workflow to account for these temporal dynamics.

Guided by the temporal gelation kinetics of the gelatin-based bioink (**Figure 1e**), we developed a precision-controlled bioprinting protocol to ensure structural consistency across bioprinting batches. Cells and bioinks were initially combined at 37°C to achieve a uniformly distributed single-cell suspension. To transition the ink into a printable state without inducing thermal shock or compromising cellular viability, the mixtures were rapidly pre-cooled in a 4°C chiller bath. We identified a critical “priming” window of 7 minutes, after which the bioink attained the necessary viscoelasticity to be extruded as continuous, high-resolution filaments. Critically, to stabilize the bioink in this optimal gelated state for the duration of the printing session (∼5 min), the printhead temperature was subsequently maintained at 21°C (**Figure 1e (2)**). Deviations from this setpoint compromised printing fidelity. Holding temperatures exceeding 21°C led to progressive reversed sol-gel transition, which caused the bioink to turn under-gelation (**Figure 1e (1))**, while temperatures below 21°C resulted in excessive solidification (over-gelation) and nozzle clogging (**Figure 1e (3)**). This rheology-informed thermal management strategy enabled consistent high-throughput production of printed vessels, providing the morphological uniformity essential for downstream functional assays and longitudinal perfusion studies.

The secondary TG crosslinking stabilized the gelatin and collagen type I components to provide essential bioactive ligands for cell attachment and proteolytic remodeling. However, the dense, ionically crosslinked bioinert alginate fraction created a physical barrier that hindered cellular spreading. To alleviate this constraint, the alginate network was selectively leached using Ca^2+^- and Mg^2+^-free DPBS. By day 7 of culture, encapsulated cells in the leached vessels exhibited robust, cell type specific morphologies and extensive cytoplasmic spreading throughout the 3D hydrogel volume (**Figure 1f**). This stood in contrast to the non-leached control group, where cells remained predominantly spherical. Scanning electron microscopy (SEM) and subsequent pore-size quantification revealed that alginate leaching induced a shift in pore size distribution, increasing the average pore diameter from 67±28μm in the non-leached controls to 107±37μm in the treated scaffolds, providing the structural basis for the enhanced cellular morphology apparent in the fully processed vessels (**Figure 1g-i**).

### Functional validation of the adventitial and medial layers

We characterized bioprinted arteries over a 21-day static culture period to assess viability and long-term stability. TUNEL and phalloidin staining were used to quantify and characterize cellular viability, morphology, and spatial distribution within the 3D architecture (**Figure 2a**). Cell viability measurements via TUNEL remained consistently above 80% throughout the study, with most apoptotic signal localized to the exterior and lumenal surfaces (**Figure 2b**). Quantifying the dimensions of the vessel wall and lumen across the 21-day time course revealed that while wall thickness remained stable, lumenal diameter increased from an initial mean of 366 μm to a final mean diameter of 629 μm at day 21 **(Figure 2c),** indicating architectural remodeling over time.

**Figure 2:**
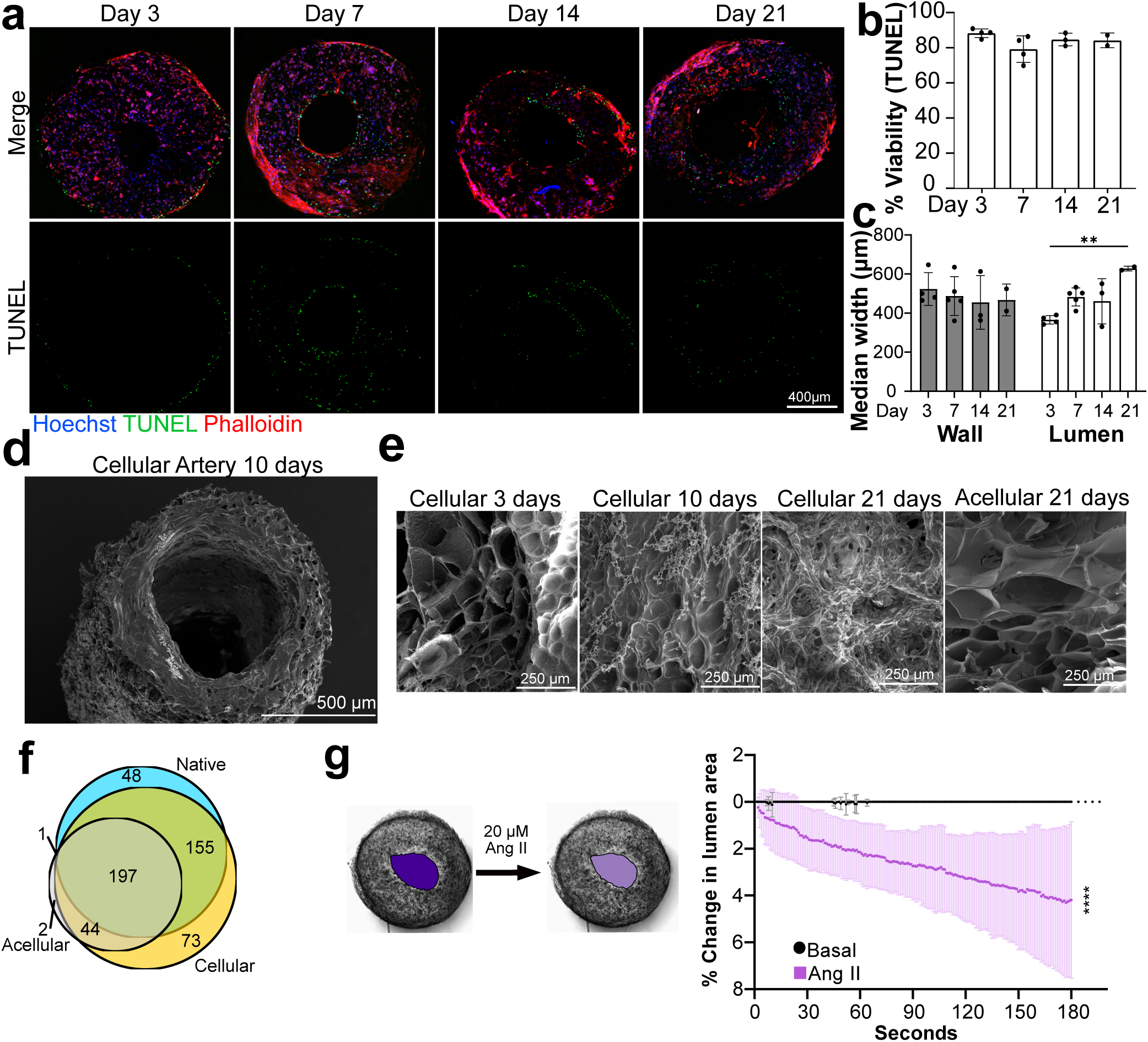
Functional adventitial and medial layers. **(a)** Representative immunofluorescence images of bioprinted vessels over 21 days in static culture stained for nuclei (Hoechst), actin (phalloidin), and dead cells (TUNEL). **(b)** Quantification of cell viability by % TUNEL-/Hoechst+ cells and vessel dimensions **(c)** at each timepoint (n=2-4 vessels, mean ± SD). **(d)** SEM image of a cellularized vessel at 21 days of culture **(e)** Representative SEM images of fibroblasts laden hydrogel on days 3, 10, and 21 of culture compared to an acellular hydrogel after 21 days of incubation in medium. **(f)** Venn diagram of matrisomal proteins present across conditions. **(g)** Quantification of percent reduction in lumenal area in response to 20pM Angiotensin II (Ang II) over a course of 180 seconds (n=9, mean ± 95% confidence interval, paired t-test, *\*\*\*p<* 0.001).

To further investigate remodeling within the bioprinted vessels, we performed scanning electron microscopy (SEM). Analysis of the arteries after two weeks of continuous perfusion confirmed that their structural integrity was well preserved, with clearly defined lumenal and wall architectures (**Figure 2d**). SEM profiling of the adventitial fibroblast-laden hydrogel revealed substantial microenvironment transformation over time (**Figure 2e)**. On day 3, the hydrogel network exhibited a porous architecture nearly identical to acellular control samples.By day 10, changes in pore and mesh size are apparent. Finally at day 21, the original hydrogel pores were no longer distinct, having been superseded by a heavily remodeled network characterized by dense fibrous depositions and organized extracellular matrix (ECM). These findings demonstrate that the encapsulated primary cells are not only viable but also actively remodeling the surrounding matrix and that the overall mechanical and structural fidelity of the artery-on-a-chip is preserved under long-term culture conditions.

To characterize this remodeling at the molecular level, we performed proteomic profiling of acellular and cellularized printed arteries cultured for 10 days in serum-containing media and compared them to native human posterior tibial arteries. The NABA matrisome gene set was used to subset ECM components from the global proteome landscape (**Figure S3a).** All samples shared a substantial set of matrix proteins, indicating that the hydrogel system recapitulates many baseline components of native artery (**Figure 2f**). The 155 proteins shared between cellularized printed vessels and native artery are deposited as the printed artery matures making the printed vessel more similar to the native artery. These proteins were most related to cell adhesion, angiogenesis, and differentiation (**Table S1**). The 48 proteins that were detected only in the native artery are associated with blood coagulation and innate immunity, which are derived from components not present in our system. Together, these findings demonstrate that cellular mediated ECM remodeling shifts the matrix proteome toward a native artery-like environment.

While fibroblasts in the adventitial layer are most associated with maintaining the ECM, smooth muscle cells of the medial layer are functionally responsible for vasoconstriction. Vasoconstriction requires smooth muscle cells that are in contractile state and are correctly oriented relative to the vessel lumen. To evaluate the physiological functionality of the vascular smooth muscle cells (VSMCs) within the bioprinted vessels, we performed vasoconstriction analysis. At basal conditions, the lumenal area of printed artery sections did not detectably change (**Figure 2h**). However, following treatment with 20 μM angiotensin II, artery lumenal area reduced by a mean of 6% after 180 seconds (**Video S2**). This reduction in cross-sectional area demonstrates the contractile capacity of the encapsulated VSMCs and confirms the successful integration of a functional medial layer within the biofabricated matrix.

### Generation and functional validation of the intimal layer

Generating a confluent endothelial monolayer across the entire lumenal surface required optimized cell-seeding to counteract gravitational settling. We implemented a rotational “flipping” strategy during the seeding process to ensure a more uniform distribution of endothelial cells (**Figure 3a**). Quantitative analysis of lumenal coverage on day 3 post-seeding showed that the bottom section maintained high confluency regardless of the method, with 93±3% for the flipping strategy compared to 89 ± 4% for static seeding. However, the top of the lumen had significantly lower coverage and was more variable without device flipping (64 ± 14%) than those devices that were flipped during EC seeding (76 ± 6%) (**Figure 3b**). At day 7 post-seeding, immunofluorescence imaging confirmed the formation of a continuous and robust endothelial layer **(Figure 3c)**. Both longitudinal and cross-sectional views demonstrated that the ECs achieved full confluency, successfully recapitulating the structural organization of the native vascular intima.

**Figure 3:**
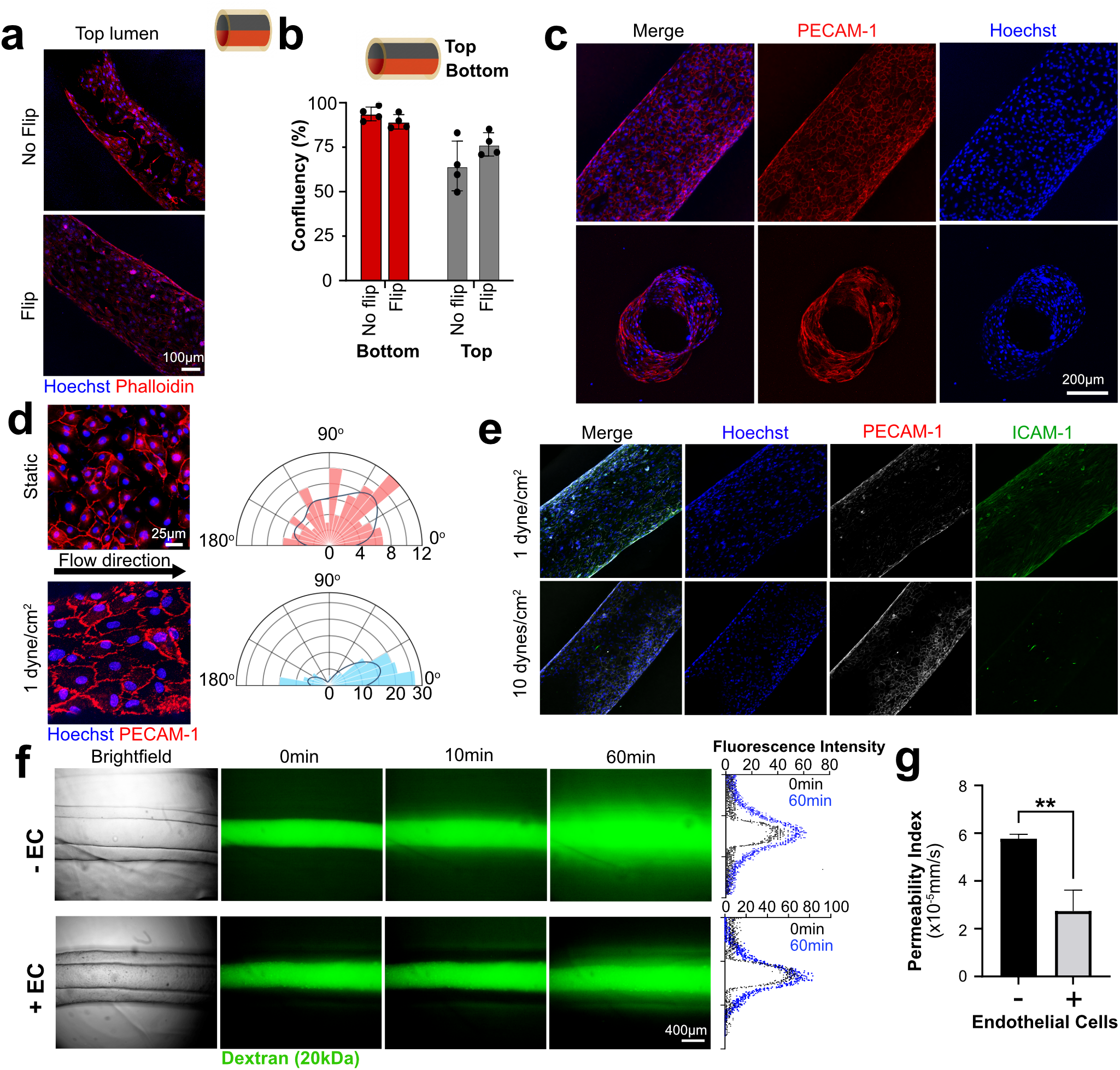
Functional validation of intimal layer. **(a)** Actin (phalloidin) staining of top lumen surface after 3 days of endothelial cell seeding, with and without flipping the device during the seeding process, **(b)** Confluence quantification of top and bottom lumen surfaces for vessels seeded with and without a 180° flipping step (n=4, mean ± SD, two-way ANOVA, * p<0.05, **p<0.01). **(c)** Representative immunofluorescence z-stack images of endothelialized lumen with flipping method after 7 days with perfusion, **(d)** Endothelial cell morphology and alignment under perfusion for 24 hours. Rose diagrams illustrate the distribution of endothelial cells alignments with the dark solid lines representing the Kernel Density Estimation (KDE). **(e)** Representative immunofluorescence images of arterial endothelial cells subjected to low (1 dyne/cm^2^) or high (10 dynes/cm^2^) shear stress for 72 hours, **(f)** Brightfield and fluorescence images showing the artery at Omin, 10min, and 60min after 20kDa FITC-dextran perfusion of bare and endothelialized vessels. The fluorescence intensity profile on the right shows the FITC intensity in a cross section through the image, **(g)** Quantified diffusional permeability index of bare and endothelialized artery (n=4, mean ± SD t-test, *\*\*\* p<* 0.001).

To verify that the endothelial layer dynamically responds to fluid mechanics, we evaluated morphological adaptation to shear stress by characterizing endothelial cell orientation and Intercellular Adhesion Molecule 1 ICAM-1 expression. Under static conditions, the ECs exhibited a stochastic orientation, with their longitudinal axes distributed uniformly across the 0°-180° range (**Figure 3d**). Conversely, introducing flow induced cell elongation and alignment parallel to the direction of flow. Quantitative analysis confirmed this transition, showing a preferential alignment within the 0° −30° and 150° −180° intervals. Investigating endothelial ICAM-1 expression revealed a non-linear response to shear stress magnitude. Under basal, static conditions, ICAM-1 expression remained minimal (**Figure S4a, b**). Low shear stress (1 dynes/cm^2^) stimulated a significant upregulation of ICAM-1 expression; however, ICAM-1 fluorescent signal returned to baseline levels at 10 dynes/cm^2^ **(Figure 3e)**. This expression pattern shows that while sub-physiological flow upregulates ICAM-1, elevating the shear stress to 10 dynes/cm^2^ suppresses this activation and restores the marker to basal levels reflecting large vessel versus capillary flow^24^.

We tested endothelial barrier integrity by measuring 20 kDa FITC-dextran diffusion kinetics. We observed rapid and extensive transmural diffusion into the medial wall and the surrounding matrix in acellular bare models immediately following perfusion (**Figure 3f**). In contrast, endothelialized vessels demonstrated significant dextran retention within the lumen, which we quantified with fluorescence intensity profiles through the arterial cross-section. The diffusional permeability coefficient (P_d_) for acellular models was 5.77×10^−5^±1.91×10^−6^ cm/s, whereas the presence of an endothelial monolayer reduced the P_d_ 52% to 2.74×10^−5^ ± 8.75×10^−6^ cm/s (p < 0.01) (**Figure 3g**). Together, the cellular alignment and ICAM presentation in response to shear and barrier activity support that the ECs in the vessel form a functional intimal layer.

### TNFα induces inflammatory responses in the printed artery that are rescued by anti-inflammatories

Having generated a functional and viable vessel with correct cellular organization (**Figure 4a, b**), we next aimed to deploy the vessel in disease modeling by introducing TNFα as an inflammatory stimulus. We perfused the printed vessels with control or 12.5 ng/mL TNFα containing media for 6 days with recirculation **(Figure 4c)**. Conditioned media was sampled daily to monitor viability and response dynamics over time and LDH release measurements showed that this stimulus did not significantly decrease cell viability **(Figure 4d).** Analyzing TNFα levels over time indicated that sustained TNFα levels persisted throughout the experiment following a single stimulation **(Figure 4e)**. Notably, TNFα concentrations exceeded those initially introduced into the perfusion system and remained detectable, consistent with sustained TNFα presence and ongoing inflammatory signaling within the perfused vessels.

**Figure 4:**
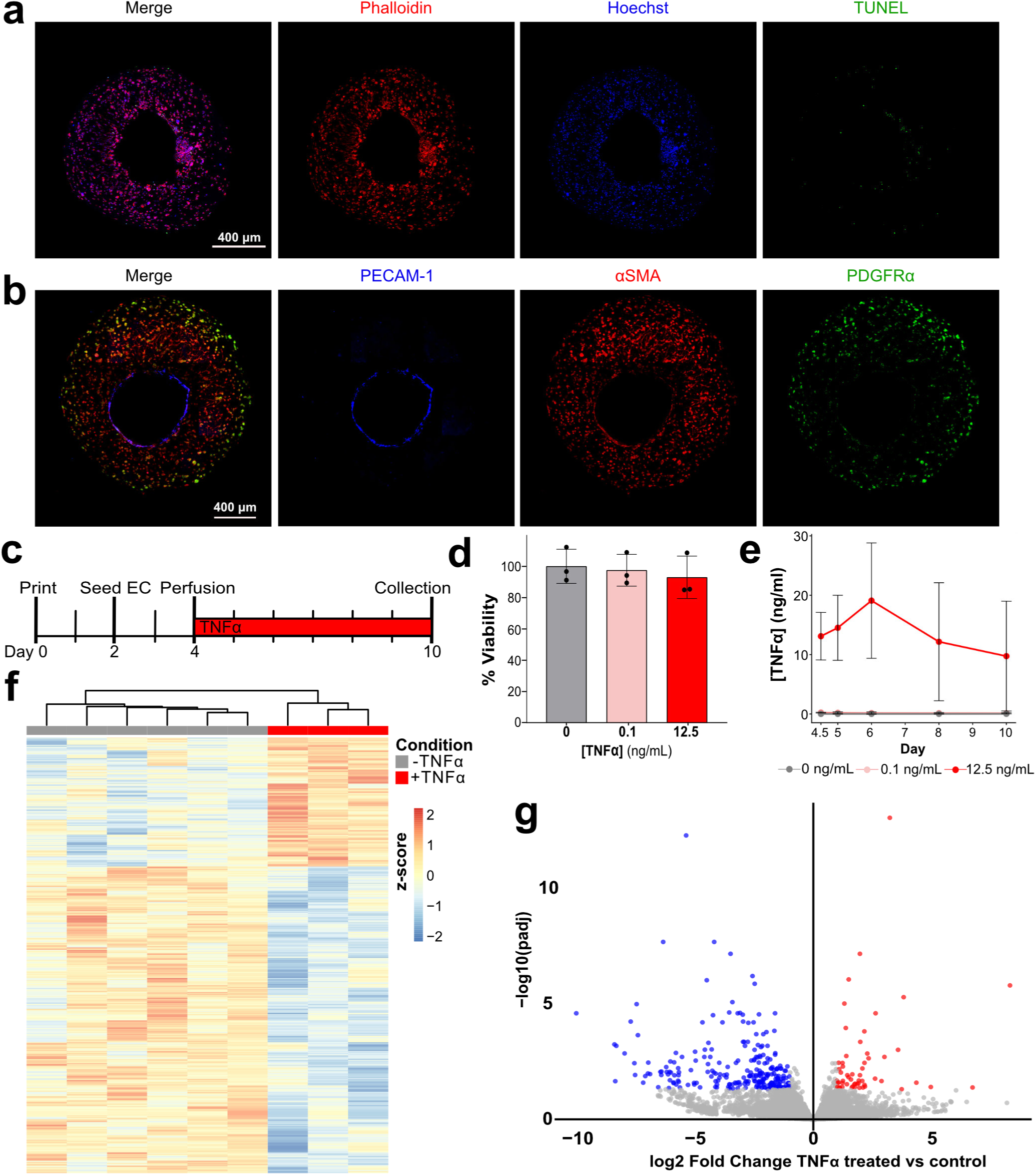
TNFα induces inflammatory response in the engineered tri-layered vessel. **(a)** Representative fluorescence images of perfused 3D vessels stained for actin (phalloidin), nuclei (Hoechst), and dead cells (TUNEL). **(b)** Immunofluorescnce of endothelial cells (PECAM), smooth muscle cells (aSMA), and fibroblasts (PDGFRa). **(c)** Experimental timeline of 3D vessel printing, endothelial seeding, perfusion, TNFα stimulation, and sample collection, **(d)** Cell viability measured by LDH assay following TNFα treatment (0, 0.1, and 12.5 ng/mL TNFα; mean ± SEM). **(e)** Perfusate TNFα levels over time in control and TNFα-treated vessels (mean ± SD). **(f)** Heatmap of row-wise z-scored variance-stabilized gene expression, showing hierarchical clustering with TNFα treatment, **(g)** Volcano plot of differential gene expression in TNFα-treated (n=3) versus untreated samples (n=6), with significantly upregulated (red) and downregulated (blue) genes indicated (adjusted p < 0.05, |log2FC| > 1).

Transcriptomic profiling revealed distinct gene expression changes associated with TNFα exposure. Hierarchical sample clustering separated TNFα-treated samples from controls, indicating a clear transcriptional response to cytokine stimulation **(Figure 4f)**. Notable transcripts upregulated in TNFα-treated samples were inflammatory mediators including *IL1B*, while downregulated transcripts were more broadly associated with proliferation and ECM remodeling (**Figure 4g, Figure S5a-b, Table S2**). This data collectively demonstrates that the printed vessels maintain organized cellular structure and viability, respond dynamically to inflammatory stimulation, and exhibit coordinated functional and transcriptional changes upon TNFα exposure.

To interrogate whether anti-inflammatory therapeutics could attenuate TNFα-induced inflammation in the engineered vessels, we evaluated secreted factors following TNFα stimulation in the presence or absence of pharmacological inhibitors. Samples were matured and perfused with either control or media containing 12.5 ng/mL TNFα to establish an inflammatory environment. A single dose of an anti-inflammatory agent, ABT-317 (a small molecule JAK inhibitor) or Infliximab (an anti-TNFα antibody) was introduced 24 hours post-TNFα insult to test therapeutic modulation of existing inflammation **(Figure 5a)**. TNFα treatment induced IL-6 accumulation over time relative to untreated controls, indicating sustained inflammatory activation within the perfused vessels **(Figure 5b)**. This elevation persisted throughout the exposure period, consistent with prolonged inflammatory signaling. To enable quantitative comparison across treatment conditions, the area under the curve (AUC) was calculated from the IL-6 time-course data. This integrated analysis demonstrated that IL-6 production was significantly reduced by both TNFα neutralization (Infliximab) and downstream pathway inhibition via JAK–STAT blockade (ABT-317) (**Figure 5c**), indicating that TNFα-driven inflammatory signaling is active in this system and can be effectively attenuated through anti-inflammatory intervention targeting either the cytokine itself or its downstream signaling pathways.

**Figure 5:**
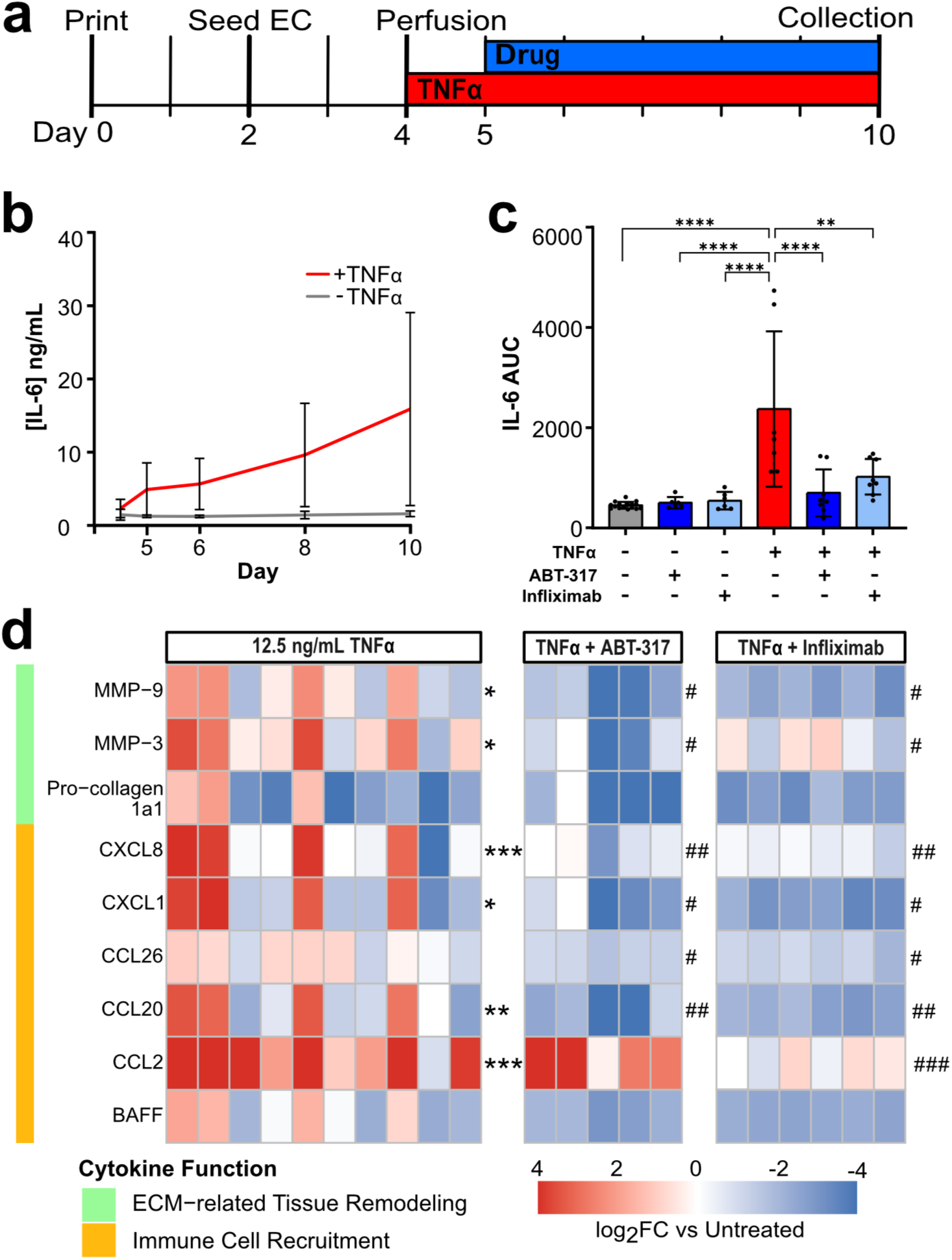
Anti-inflammatory treatments suppress vessel inflammation, **(a)** Experimental timeline demonstrating TNFα stimulation followed by anti-inflammatory treatment (JAK inhibitor ABT-317, anti-TNFα infliximab), **(b)** Temporal IL-6 levels in perfusate from control and TNFα-treated vessels and **(c)** corresponding calculated area under the curve for vessels ± TNFα and ± anti-inflammatory treatments (1 pM ABT-317 or 10 ug/mL infliximab) (n=5-10, mean ± SD, one-way ANOVA). **(d)** Heatmap depicting log2FC changes in the AUC of secreted factors relative to untreated controls (n=14) after TNFα stimulation (n=10) and drug intervention (n=5-6). Statistical significance by one-way ANOVA is denoted as * *p* < 0.05, *\*\* p<* 0.01, and *** *p <* 0.001 versus untreated controls, and *# p* < 0.05, *##p* < 0.01, ###*p <* 0.001 versus TNFα.

To study these interactions beyond IL-6, multiplex Luminex analysis was performed using a custom-designed panel of inflammatory and remodeling-associated factors selected to capture key targets downstream of JAK–STAT signaling that are implicated in stromal immune cell recruitment and ECM remodeling. TNFα stimulation resulted in coordinated significant increases in cytokines and chemokines associated with extracellular matrix remodeling and immune cell recruitment, including MMP-9, MMP-3, CXCL8, CXCL1, CCL20, and CCL2, while treatment with either ABT-317 or Infliximab significantly attenuated this response **(Figure 5d, S6)**, highlighting both the induction and effective suppression of this inflammatory and remodeling response in this system.

## DISCUSSION

Recent engineered arteries that incorporate multiple vascular cell types have established bioprinting as a feasible method to generate multi-layered vessels for engraftment^25–29^ or *in vitro* experimentation. Many *in vitro* arterial models primarily examine the effects of mechanical stimulation on the artery especially in relation to early markers of atherosclerosis^30–35^. Fewer models are deployed to investigate the impact of cytokines or conditioned media^36–39^. While these models generated valuable technical progress and biological insight, important gaps remain including incorporating tissue specific adventitial cells, expanding experimental readouts and timelines, and streamlining support devices to reduce proteins absorption and simplify terminal sample recovery. We established a tissue engineering process to produce anatomically representative bioprinted vessels from primary human arterial cells to model vascular inflammation. Our findings emphasize the importance of quality control steps to incorporate primary human arterial cells, establish narrow parameters to reproducibly generate bioprinted arteries, and interrogate the response of our system to TNFα insult and ability to resolve by treatment with both small molecule JAK inhibitor ABT-317 and monoclonal antibody infliximab. We have positioned our model as an accessible and flexible platform to examine both arterial response and therapeutic treatment.

Multi-material coaxial printing is a powerful technique to efficiently deposit multiple layers simultaneously which we adapted to generate tri-layered vessels. With gelatin-based hydrogels, managing the non-linear kinetics of thermal solid-gel transition is crucial to ensure structural fidelity across multiple samples. We determined that an initial thermal equilibration at 4℃ rapidly drove the cell-laden bioink into printable state and that a narrow window around 21℃ maintained optimal shear-thinning viscoelastic consistency for an extended operational time frame^27^. Failure to strictly maintain the bioink within this narrow thermal boundary results in either melting to an unprintable liquid or rapid, uncontrolled physical crosslinking that introduces severe structural variance. Attempting to correct over-gelation by re-heating the ink or under-gelation by chilling the ink necessitates an entire system reconfiguration, dramatically expanding the time that primary cells are subjected to suboptimal conditions. Consequently, this multi-stage thermal stabilization protocol provides structural uniformity and sufficient samples required for downstream multiplexed evaluations.

Sequencing and cell morphology studies demonstrate substantial heterogeneity between cells of the same type derived from different tissues or even the same tissue but different anatomical location^40–44^. Incorporating tissue specific primary cells can provide increased tissue fidelity but requires additional quality control measures. One well known challenge to incorporating primary smooth muscle cells and fibroblasts is the spectrum of cell states that these cells adopt. We characterized available cell lots prior to use and reemphasize that relying solely on manufacturer labeling or immunofluorescence is insufficient to guarantee correct cell state. By prescreening primary cells and limiting passages on hard plastic, we were able to maintain characteristic arterial anatomical arrangement for extended culture. Notably, we detected phenotypic plasticity within the 3D model over time. Indeed, in longer experiments our model may capture the adaptive cellular transitions characteristic of primary human vascular tissues^45,46^.

An essential consideration for our microphysiological blood vessel is the mechanical environment imposed by fluidic parameters. The magnitude of shear stress detected by ECs regulates ICAM-1 expression^47,48^. We found that under low-shear or static conditions ECs upregulated ICAM-1 (**Figure S3a**). Conversely, increasing the mechanical stimulus toward arterial physiological range to 10 dynes/cm^2^ suppressed ICAM-1 expression back to baseline (**Figure 3h**). This localized suppression demonstrates that the endothelium within our platform responds appropriately to mechanical cues to regulate cellular activation and maintain homeostasis^24,49^. Driven by the pulsatile flow, the vessel wall undergoes continuous cyclic stretch and recoil. This dynamic physical deformation effectively trains the smooth muscle cells via mechanical stretch, providing the structural guidance cues necessary to drive their circumferential alignment along the lumen^50^. Recapitulating concentric cellular organization mirrors the highly structured medial layer of native human arteries *in vivo* and is essential for effective vasoconstriction in our model (**Figure 2g**). Together, this stretch-trained smooth muscle cell alignment and the endothelial response to flow highlight the robust organotypic characteristics of our bioprinted vascular models.

The bioprinted artery responded to TNFα perturbation and that response was resolved by ABT-317 or infliximab treatment. While TNFα mediated endothelial barrier disruption is well established^38,51^, less attention has been paid to the secretory response and vascular cell interaction. We incorporated all three arterial cell types and stimulated with sufficient TNFα to induce a response without compromising viability. We detected significant shifts in the secretory profile of the vessel that would contribute to a sustained inflammatory response. Notably, the most significantly upregulated transcript after TNFα stimulation was *IL1B* and other transcripts that also changed are involved in IL1B response (**Table S2**). Others with various methods of cytokine insult to engineered arteries also found increased *TNFα* and *IL1B* transcripts levels^37,39^. Curious about the potential feedforward signaling, we compared the transcripts that changed with TNFα stimulation with ChIP-seq of transcription factors downstream of TNFα, IL1B, and other signaling molecules detected in the transcriptomic data (**Figure S6**). We found substantial overlap, suggesting that future work could pursue the timing and individual effects of signaling pathways activated in vascular inflammation.

Anti-inflammatory treatment reversed TNFα-induced secreted proteins as a single dose administered 24 hours after TNFα stimulation. Reducing the response to TNFα post TNF administration rather than prophylactically or concurrently with two different classes of anti-inflammatory indicates the strength of our model. Together these results support that the bioprinted vessel can model secondary vascular inflammation brought on by circulating cytokines and that secondary effects of chronic inflammation can be pursued through complex *in vitro* models. Increasing the complexity of our bioprinted vessel system in future work will expand the applicability as a complex representation of *in vivo* interactions. Cellularizing the support gel to model specific tissue types, like lung or lymphoid tissue, will facilitate studies of vessel-tissue interaction. Incorporating immune cells into the vessel both in the adventitial layer and circulating through the lumen will further expand disease states that can be appropriately modeled. Our model provides a modular and flexible organotypic human vessel for recapitulating and perturbing complex biological processes *ex vivo*.

## FIGURE LEGENDS

**Supplemental Figure S1:**
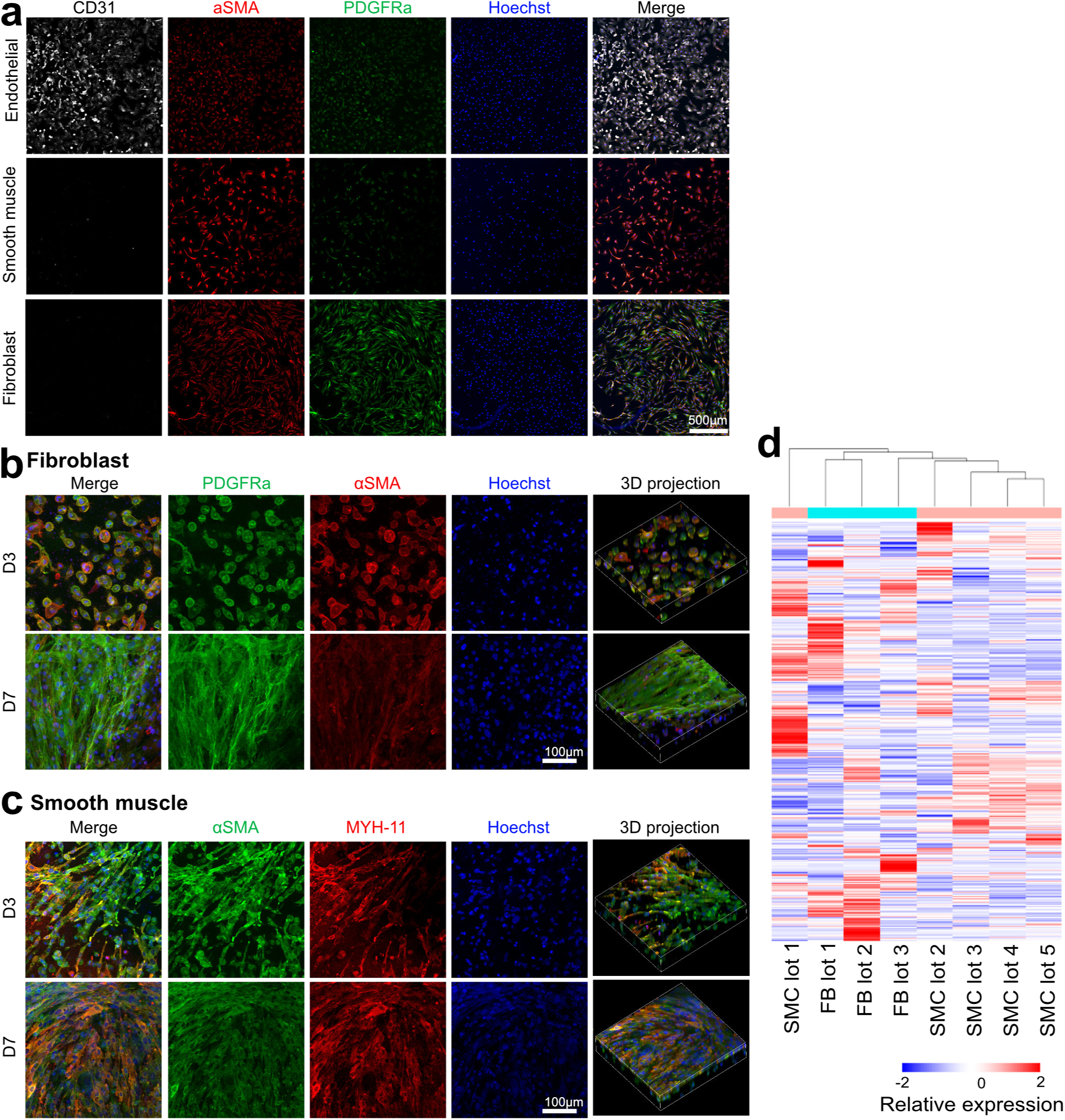
**(a)** Representative immunofluorescence images of primary human arterial endothelial cells, smooth muscle cells, and fibroblasts. Immunofluorescence images of **(b)** smooth muscle cells and **(c)** fibroblasts encapsulated in hydrogel at 3 or 7 days of 3D culture, **(d)** hierarchical clustering of top 200 transcripts of three fibroblast and 5 smooth muscle cell lots.

**Supplemental Figure S2:**
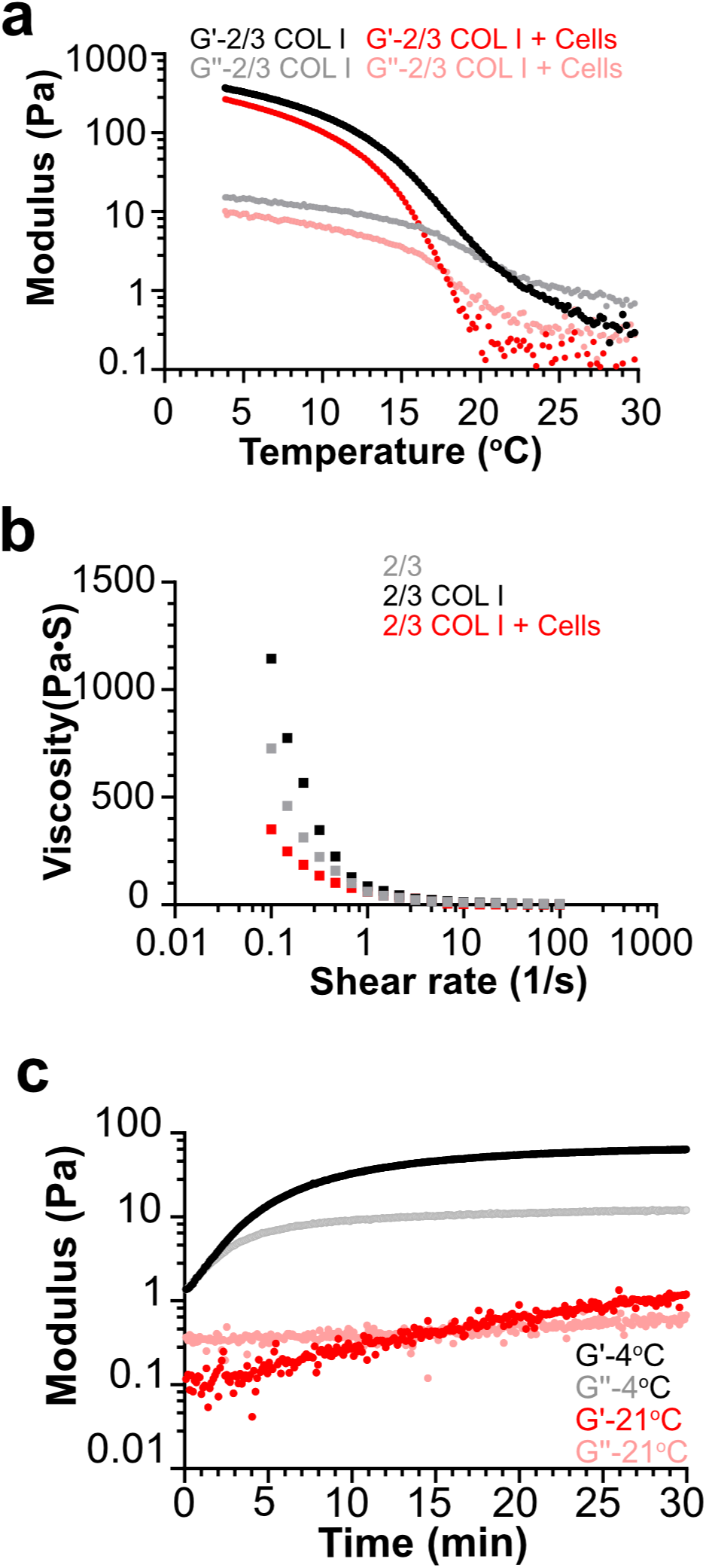
**(a)** Temperature ramp analysis of thermoresponsive gelatin-alginate-collagen based bioink with and without cells, **(b)** Shear rate ramp analysis characterizing shear thinning properties of gelatin-alginate, gelatin-alginate-collagen, and cell-laden bioinks, **(c)** Time-dependent thermo-gelation of cellular gelatin-alginate-collagen bioink at 4°C and 21°C for a period of 30 minutes.

**Supplemental Figure S3:**
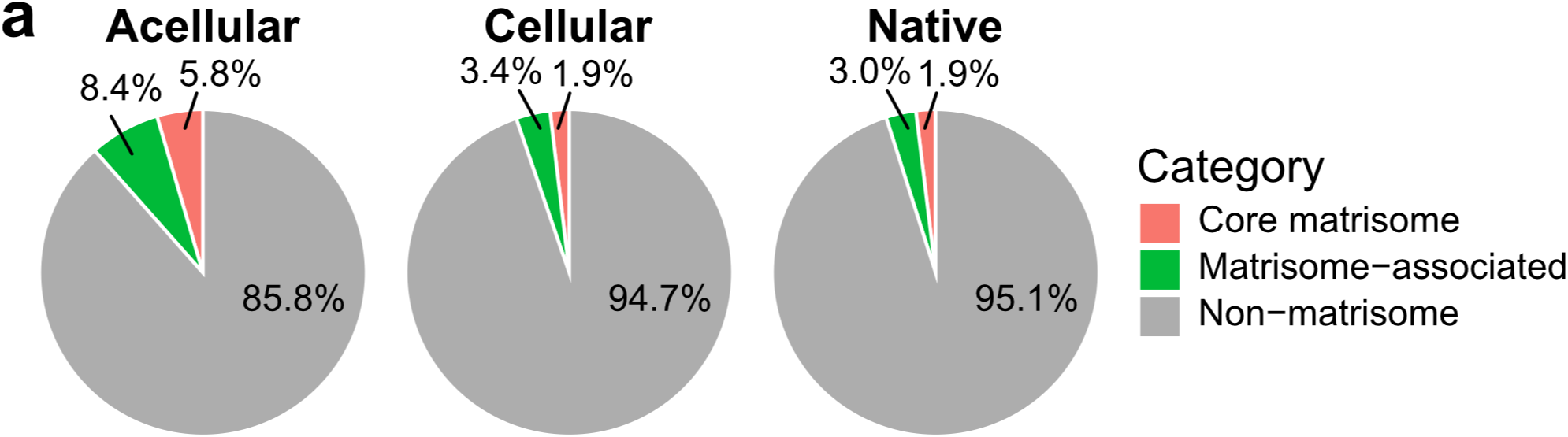
**(a)** Percentage of proteins detected in acellular printed vessels, cellular printed vessels, and native arteries classified as matrisome, matrisome-associated, or non-matrisome proteins^52^.

**Supplemental Figure S4:**
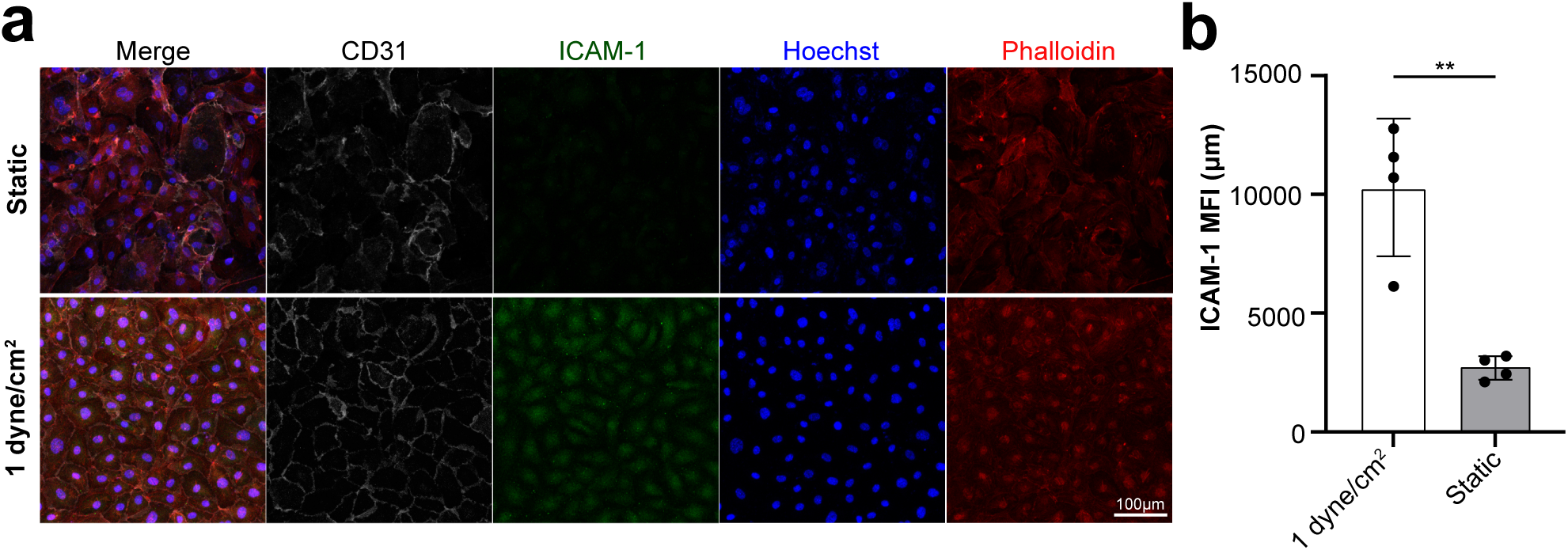
**(a)** Representative immunofluorescence images of arterial endothelial cells subjected to static culture and low shear stress conditions (1 dyne/cm^2^) for 72 hours. **(b)** Average ICAM-1 fluorescence intensity per cell under static and low shear culturing conditions (**p<0.01).

**Supplemental Figure S5:**
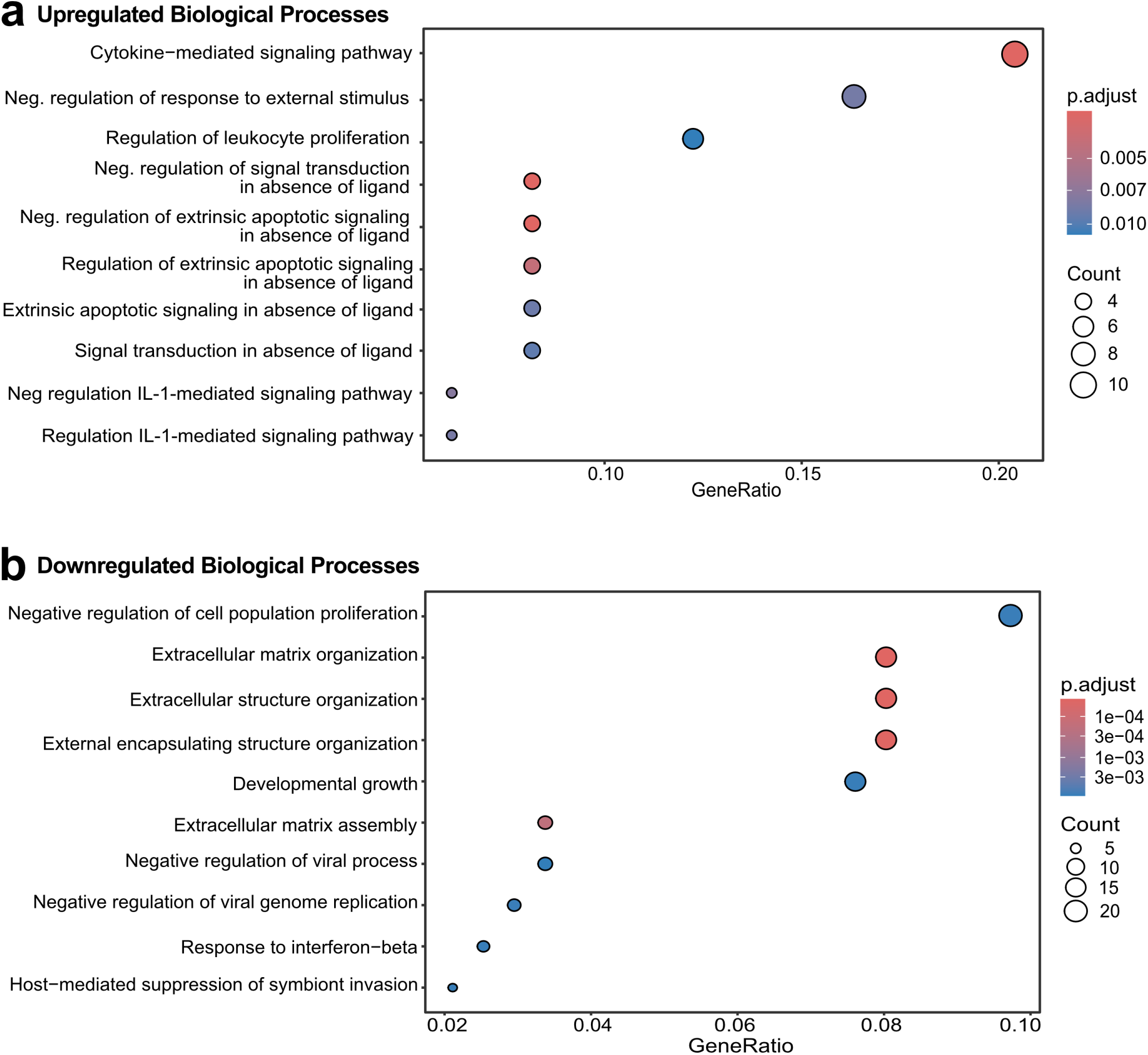
Top ten most significant GO biological process terms for the **(a)** up regulated transcripts (56) or **(b)** down regulated transcripts (255) following TNFα stimulation.

**Supplemental Figure S6:**
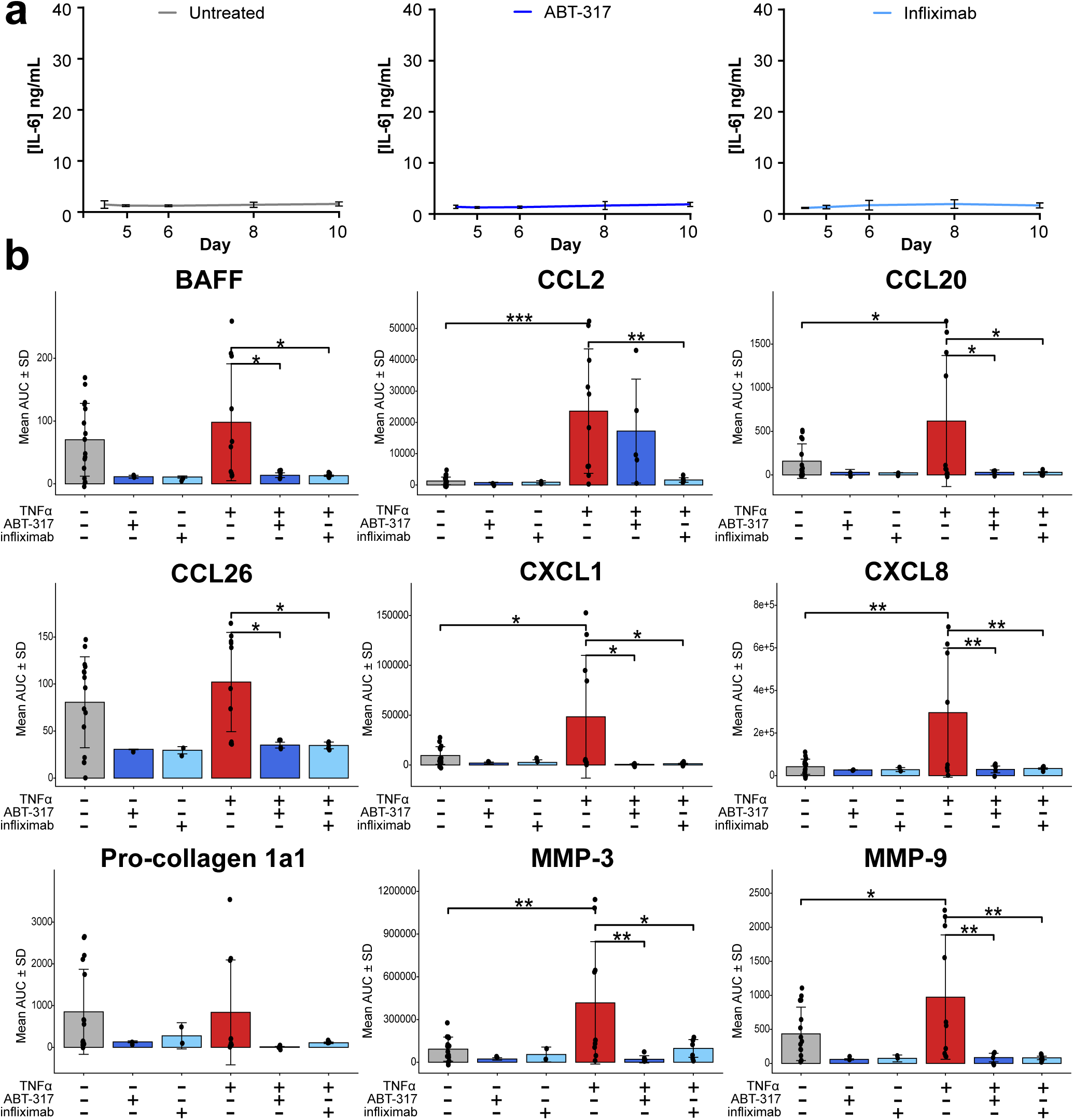
**(a)** Temporal IL-6 levels in perfusate from control, ABT-317, and infliximab treated vessels**. (b)** Individual AUC profiles for all detectable analytes measured in the multiplex Luminex panel, including BAFF, CCL2, CCL20, CCL26, CXCL1, CXCL8, Pro-collagen 1a1, MMP-3, and MMP-9. AUC values represent cumulative secretion over the experimental time course. Data corresponds to the same dataset summarized in the heatmap (Figure 5d) and are shown here as non-normalized, per-analyte profiles. Vessels were treated with TNFα (n = 10) and in combination with anti-inflammatory treatments, including 1 pM ABT-317 or 10 μg/mL Infliximab (n = 5-6). Data are presented as mean ± SD with individual datapoints. Statistical significance by one-way ANOVA is denoted as * p < 0.05, ** p < 0.01, and ***p< 0.001.

**Supplemental Video 1**: Example free fall print of a vessel into a CaCl_2_ bath.

**Supplemental Video 2:** Representative timelapse of a vessel treated with 20 μM Angiotensin II. Total time 180 seconds acquired at 1 frame per second.

## METHODS

### Cell Culture

Primary human aortic smooth muscle cells (VSMC) (Cell Systems, Cell Biologics) were cultured in Ham’s F-12K (Kaighn’s) medium (Gibco, 21127030) supplemented with 10% FBS (Gibco, A5670701), 1X Penicillin-Streptomycin (MP Biomedicals, 091670454), 2mM L-glutamine (Gibco, 25030081), 5 ng/mL insulin (Sigma Aldrich, 91077C250MG), 10 ng/mL epidermal growth factor (EGF) (R&D Systems, 236EG200), and 1 ng/mL fibroblast growth factor (FGF) (R&D Systems, 3718-FB). Primary human aortic adventitial fibroblasts (Cell Applications, Cell Biologics) were cultured in DMEM medium (Gibco, 11965118) supplemented with 10% FBS, 1X Penicillin-Streptomycin, 2mM Sodium Pyruvate (Gibco, 11360070), 2mM L-glutamine, 5 ng/mL insulin, and 1 ng/mL fibroblast growth factor (FGF). Primary human aortic endothelial cells (Cell Applications) and were cultured in Cell Biologics Complete Endothelial Cell Medium Kit (Cell Biologics, H1168). All cells were cultured at 37°C in 95% humidified air containing 5% CO_2_. Cells were used within the first ten passages. Tri-layer arterial vessels were cultured in a co-culture media made up of Ham’s F-12K (Kaighn’s) medium supplemented with 10% FBS, 1X Penicillin-Streptomycin, 2mM L-glutamine, 5 ng/mL insulin, 10 ng/mL epidermal growth factor (EGF), 1 ng/mL fibroblast growth factor (FGF), 90 μg/mL heparin (Thermo Scientific, AAA1619803), and 1 ng/mL vascular endothelial growth factor (VEGF) (R&D Systems, BT-VEGF).

### Rheology

The tested formulations include an acellular control composed of 2% (w/v) alginate, 3% (w/v) gelatin, and 0.5 mg/mL Type I collagen, as well as a cellular bioink consisting of the same hydrogel matrix encapsulated with a cell density of 1×10^7^ cells/ml. The rheological behaviors of the bioinks were measured using a TA Discovery HR 20 equipped with a cone and plate geometry. Thermal gelation properties were assessed using temperature ramp from 30 °C to 4 °C at a step size of 2°C /min. The complex modulus was measured according to shear rate ramp from 0.1 1/s to 100 1/s at 4°C for 15 minutes to evaluate shear thinning characteristics. A time sweep analysis was performed with an initial 1-minute 37°C incubation followed by a 30-minute test at 4°C and 21°C to confirm gelation properties over time at relevant workflow temperatures. For temperature tuning on gelation properties, the samples were first equilibrated at 37°C for 1 minute, followed by rapid cooling at 4°C for 7 minutes to induce gelation. Subsequently, the temperature was adjusted to specific increments (19°C, 20°C, 21°C, and 22°C) to monitor the stability and phase transition status of the bioink.

### 3D Bioprinting of Artery Constructs

Artery constructs were printed using an Allevi 3 extrusion bioprinter (3D Systems) with a composite alginate–gelatin–collagen bioink. The base bioink was prepared by dissolving 2.5%(w/v) alginic acid sodium salt, very low viscosity (Thermo Fisher, A1856522) in PBS, pH 7.4 (Gibco, 10010023) at 50°C, followed by the addition of 3.75%(w/v) porcine gelatin (300 g bloom) (Sigma Aldrich, G1890100). The solution was sterilized by heating to 70°C for 30 minutes, cooling to 4°C, and repeating the cycle two more times. Type I collagen solution was neutralized by combining VitroCol® Type I Human Collagen Solution (3.1 mg/mL; Advanced Biomatrix, 5007) with 0.1 M sodium hydroxide (Fisher Scientific, SS267) and 10X PBS (Fisher, BP2944100) at an 8:1:1 ratio, yielding a final concentration of 2.48 mg/mL. Prior to cell incorporation, the sterile alginate-gelatin bioink was liquified at 37°C. Cells were gently resuspended in four parts base alginate-gelatin ink and one part neutralized collagen I to achieve a final concentration of 1×10^7^ cells/mL, with a final composition of 2% alginate, 3% gelatin, and 0.5 mg/mL collagen I. The cellularized bioink was loaded into a 5mL Allevi plastic syringe and placed in chiller (Launa) set to 4°C for 7 minutes or until bioink reached a gel like consistency.

### Printing Parameters

Bioink was immediately transferred fort eh chiller to temperature-controlled print heads and maintained at 21°C for the duration of printing. Vessels were free fall extruded from the custom nozzle (inner 22 G, middle 15G, Outer 11G) into a bath containing an initial crosslinking solution (200 mM CaCl_2_ (Thermo Fisher, L13191.30), 6 U/mL Transglutaminase, 5% Fetal Bovine Serum (Gibco, A5670701) and 25 mM HEPES (Gibco, 15630106) in high purity deionized (DI water) for alginate crosslinking. Vessels were printed in 100 second bursts.

### Post-Print Processing

Following 5-minute incubation, the initial crosslinking solution was removed, and the arteries were briefly rinsed with a solution of 20 mM CaCl_2_ and 25 mM HEPES in high purity DI water. Crosslinking of gelatin and collagen I was then performed by incubating the arteries for 2 hours at 37°C in co-culture media supplemented with 6 U/mL Transglutaminase^27^ and 20 mM CaCl_2._ Arteries were subsequently cannulated onto prepared devices and 10% low melting point agarose (IBI Scientific, IB70056) in high purity DI water was applied around the cannulation nozzles to secure the vessels. To remove residual calcium, arteries were washed with Calcium Magnesium free DPBS (pH 7.4) (Gibco, 10010023). Alginate was leached from the arteries by adding 1 mL of DPBS (pH 7.4) to the center chamber of each device and incubated at 37℃ for 20 min based on previously reported protocol^53^. DPBS was finally replaced with co-culture media and maintained at 37°C in 95% humidified air containing 5% CO_2_ until endothelial cell seeding.

### Device Fabrication

The artery housing units were 3D printed using BioMed White Resin (Formlabs, RS-F2-BMWH-01) on the Form 3 Printer (Formlabs). All devices were washed and sterilized based on manufacturer recommended protocol. A polypropylene nozzle with luer-lock connection (1-1/2” Long 22 Gauge Needle) (McMaster-Carr, 6934A112) was threaded through the side ports and glued in place using LOCTITE 5861 (Henkel, 518485). KWIK-SIL low toxicity silicone bio-adhesive/sealant (WPI, KWIK-SIL) was added around the perimeter of the bottom of the device before securing a rectangular coverslip (24 mm x 60 mm) in place. The glue was allowed to cure overnight. The housing units were then sterilized by submersion 70% IPA for 5 minutes and UV exposure for 30 minutes on both the top and bottom sides.

### Endothelial Cell Seeding

5% low melting point agarose (IBI Scientific, IB70056) was dissolved in Ham’s F-12K (Kaighn’s) Medium (Gibco, 21127030) at 37°C. Approximately 1 mL was added to the center chamber of each device and allowed to solidify for 1 minute. Endothelial cells (2×10^7^/mL) were slowly added to the lumen of the artery via the cannulation ports. Uniform lumenal coverage was achieved by axially inversion (180°) of the device with gentle mechanical agitation, repeated 3 times at 15-minute intervals. Co-culture media was subsequently added to the device, and arteries were maintained at 37°C in 95% humidified air with 5% CO_2_ until ready for perfusion.

### Preparing Devices for Perfusion

Medium was thoroughly aspirated off the top of the device to ensure the top surface was completely dry. KWIK-SIL low toxicity silicone bio-adhesive/sealant (World Precision Instrument, KWIK-SIL) was added around the perimeter of the top surface of the device before firmly securing cover glass (24×50 #1 thick, Knittel Glass) on top and allowed to cure for 1.5 hours in the cell culture incubator before being subjected to perfusion.

### Perfusion System Assembly

A closed-loop perfusion system was assembled using commercially available plastic fittings and luer-lock components. A peristaltic pump (Golander, BT100S-1) was used to generate flow of 40 μl/min. A 50 mL falcon centrifuge tube (Corning, 352070) served as a media reservoir and was fitted with a custom 3D-printed screw cap with three GL14 ports (grabcad.com/library/50ml-falcon-lid-multi-bola-gl14-1). The cap was fabricated using BioMed White Resin (Formlabs, RS-F2-BMWH-01) on the Form 3 Printer (Formlabs) in accordance with the resin manufacturer’s instructions. GL14 screwcaps (DWK, 292400806) with 5/32” holes drilled into the tops were fitted onto each of the GL14 ports to allow tubing to pass through. Custom O-ring fittings used to create a seal between the screw caps and the GL14 ports were made by cutting 4 mm sections from DURAN® GL 14 Blanking Screw Cap and using a 3mm biopsy punch (Bianco Brothers, BB-BP-03) to create a hole in the center. Three Sani-Tech® STHT®-C Silicone Tubing (Saint-Gobain, STHT-C-062-1) lines were passed through the caps and O-ring fittings to enable inflow, outflow, and gas exchange with the media reservoir.

Sani-Tech® STHT®-C Silicone Tubing (Saint-Gobain, STHT-C-062-1) was used as the primary tubing for the system excluding a 20 cm section of PharMed® BPT which was required for compatibility with the peristaltic pump. Tubing sections were connected via plastic barbed straight tube fitting (McMaster-Carr, 5047K71). The peristaltic pump was positioned downstream of the media reservoir outflow and upstream of the device. The device which was connected to the tubing lines using luer lock tube couplings (barbed for 1/16” tube ID) (McMaster-Carr, 51525K281). An inline sampling and injection port (Ibidi, 10820) was inserted between the device outlet and media reservoir inflow using luer lock tube couplings (McMaster-Carr, 51525K281 & 51525K271). A one-way syringe filter (0.22 μm pore size) (fisher, FIS09-720-511) was connected to the third port on the media reservoir using a luer lock tube coupling to permit gas exchange.

For reuse, tubing was flushed with 70% isopropyl alcohol followed by high purity deionized (DI) water for 5 minutes. The tubing was then dried with nitrogen gas and autoclaved to ensure sterility.

### Vessel stimulation

Vessels were perfused with co-culture medium alone or with co-culture medium supplemented with 12.5 ng/mL recombinant human TNFα (R&D Systems). For drug treatment a single dose of 1000nM ABT-317 or 10 μg/mL Infliximab (Med Chem Express) was added to the media reservoir and maintained throughout the remainder of the experiment.

### A closed-loop Immunofluorescence and fluorescent staining

3D samples were fixed with 4% paraformaldehyde for 1 hour, followed by a 20 minute PBS wash, then incubated with blocking and permeabilization buffer (3% BSA, 0.1% Triton X-100 in PBS) for 1 hour at room temperature. Primary antibodies: anti-CD31 (1:25) (BD Biosciences, 550389), anti-αSMA (1:500) (Invitrogen, 14976082), anti-PDGFRα (1:250) (abcam, ab203491), anti-MYH11 (1:200) (Invitrogen, MA5-42845) and/or anti-ICAM1 (1:100) (proteintech, 10831-1-AP) were diluted in the block/permeabilization buffer and incubated with the samples at 4°C overnight. After a PBS wash, the samples were incubated with the secondary antibodies: Alexa Fluor 488 (, 555, and/or 647 (1:500), and any additional fluorescent dyes: Hoechst 33342 (1:1000), Alexa Fluor 555-conjugated phalloidin (1:400), at room temperature for 4 hours. 2D samples were stained using the same processes with shortened incubation steps–15-minute fixation, 30-minute block/permeabilization, 1-hour primary incubation, and 30-minute secondary incubation.

### Viability

Sample viability was evaluated by microscopy or colorimetric assay. By microscopy, live samples were processed with Invitrogen™ LIVE/DEAD™ Viability/Cytotoxicity Kit for mammalian cells (Fisher, L3224) following the manufacturer’s instructions. Fixed samples were stained with the Novus Biologicals™ OneStep TUNEL Apoptosis Kit, Green, 488 (Fisher, NB170654), with minor modifications for 3D constructs. For TUNEL, a 1-3mm artery segment was cut and fixed with 4% paraformaldehyde for 1 hour followed by blocking and permeabilization with 0.1% Triton X-100 in 3% BSA, for 1 hour at room temperature. For both methods, images were analyzed using FĲI/ImageJ (NIH, USA), and cell viability was determined as the ratio of live cells to the total number of cells. Conditioned media samples were analyzed by colormetric samples with CytoTox 96® Non-Radioactive Cytotoxicity Assay (Promega, G1780) following the manufacturer’s instructions.

### Scanning Electron Microscopy

Hydrogel vessels were flash-frozen in liquid nitrogen and lyophilized to preserve the porous architecture of the polymer network. The freeze-dried matrices were mechanically fractured to expose internal cross-sectional pore structures and mounted onto metal stubs using conductive carbon adhesive tape. To introduce conductivity and prevent surface charging during analysis, the samples were sputter-coated with a 2 nm layer of gold nanoparticles. Microstructural characterization and pore morphology visualization were conducted using a Teneo field-emission scanning electron microscope (Teneo LVSEM, Thermo Fisher Scientific). Imaging was executed in secondary electron (SE) mode utilizing an Everhart-Thornley detector (ETD) with an accelerating high voltage (HV) of 5.00 kV and a probe current of 0.4 nA.

### RNA isolation

RNA was isolated from 5mm segments of printed vessels recovered from the support hydrogel after 6 days of perfusion or 20 μL hydrogel drops after 7 days of static culture by digestion in 12.5 mg/mL collagenase II (Sigma-Aldrich, C2-28-100MG) at 37°C for 20 minutes to isolate the cells from the hydrogel. Cells were pelleted by centrifugation for 5 minutes at 300xg, and RNA was extracted using the Qiagen RNeasy Plus Mini Kit (Qiagen, Cat No #74136) according to the manufacturer’s protocol.

### q-RTPCR

To establish a relevant TNFα concentration we generated dose-response curves for each cell type in 2D culture as measured by qPCR (**Figure S7a-c**). Accounting for the different responses of each cell type and transcript, we determined that 12.5 ng/mL TNFα would yield a reliable response in all cells included in the model. Isolated RNA was reverse transcribed to cDNA with Invitrogen Superscript IV. 20ng of cDNA was input into SYBRgreen based quantitative real time PCR reactions with primers that hd been validated for primer efficiency. Results were analyzed by -log2ΔΔCT.

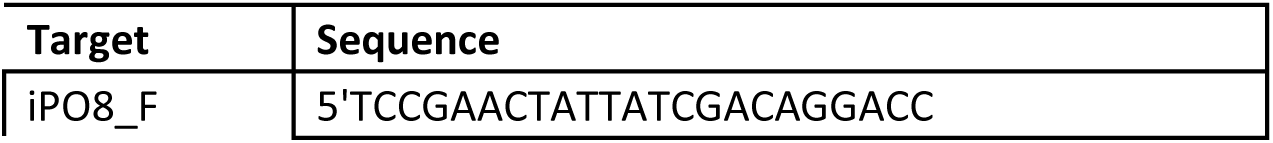
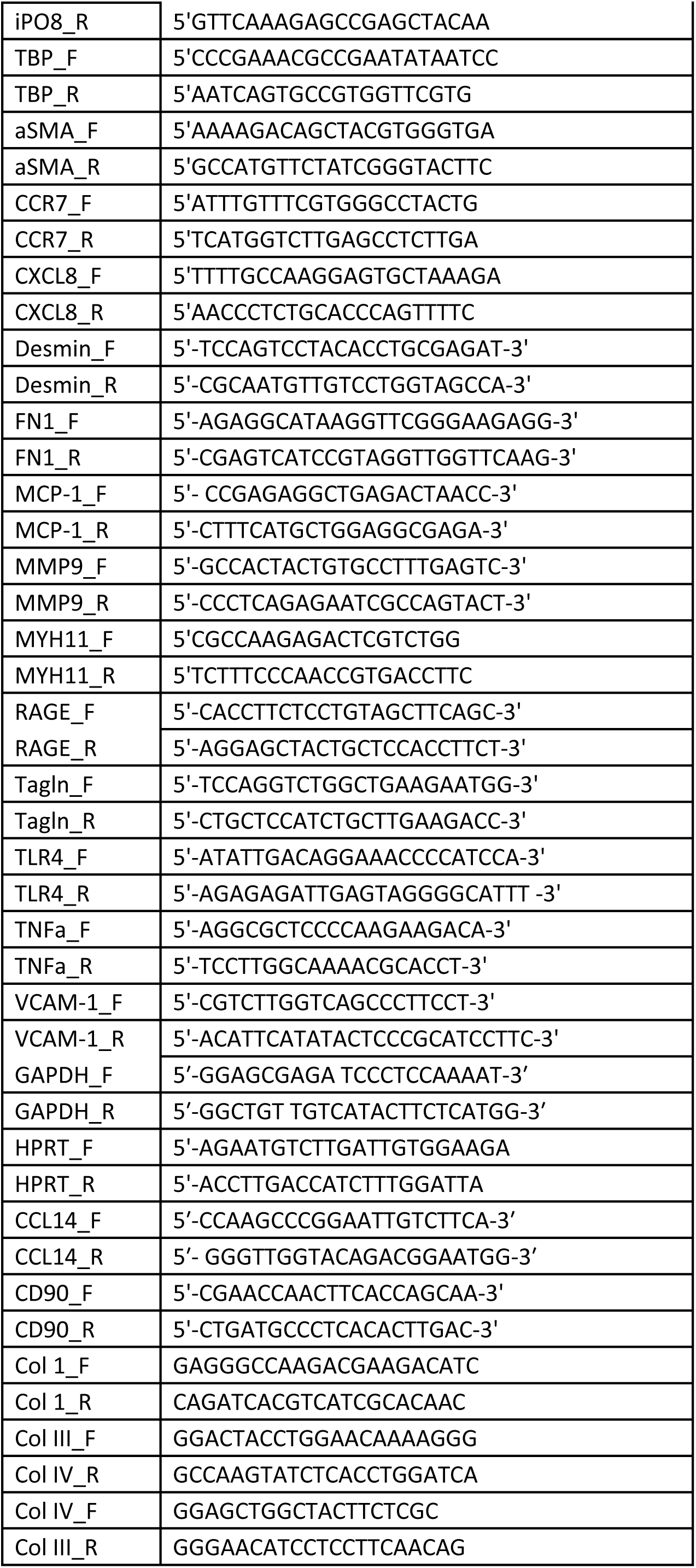
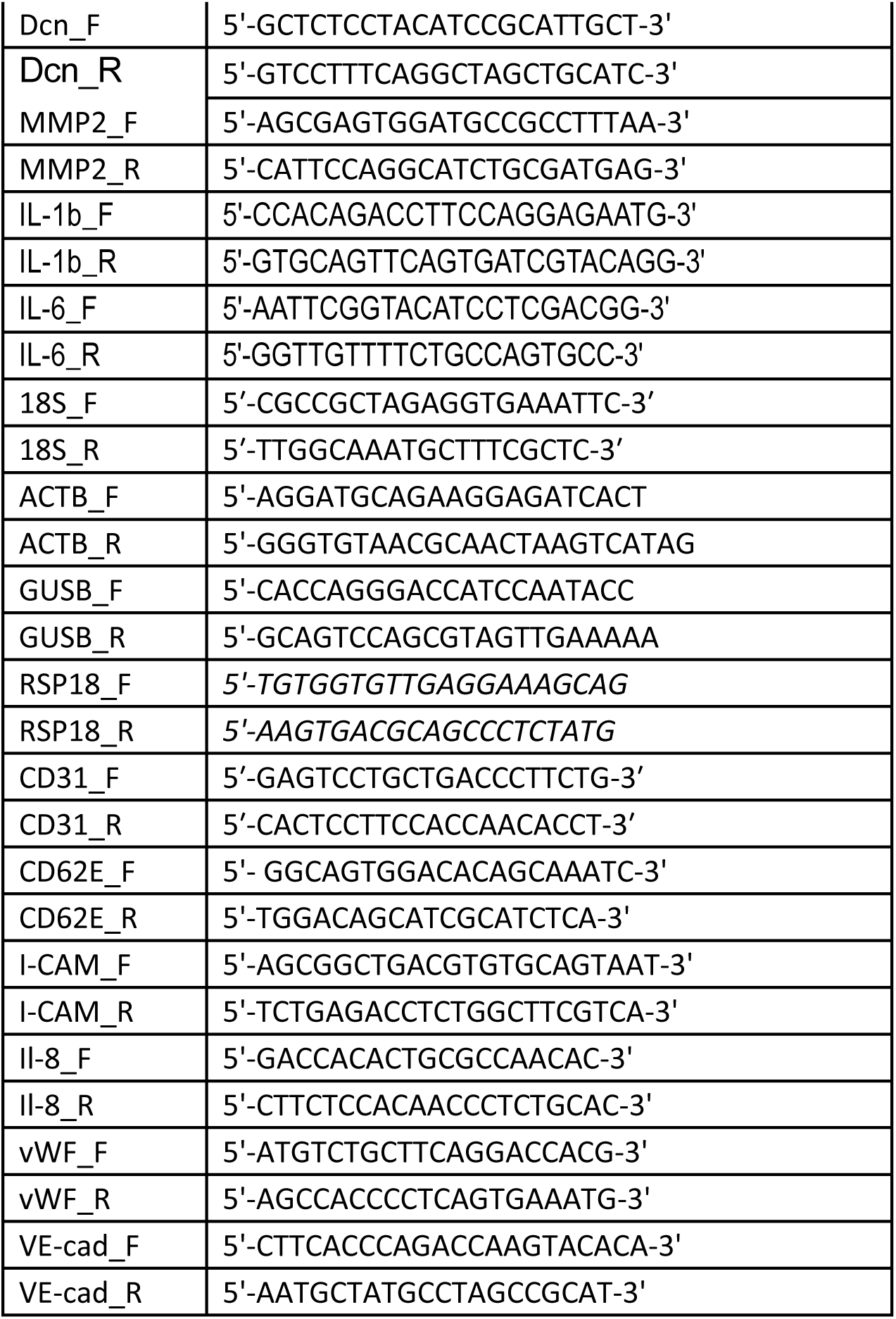

### Transcriptomics

Isolated RNA was sequenced by Novogene using standard methods outlined below.

#### Library preparation for Transcriptome sequencing

Non strand specific library Messenger RNA was purified from total RNA using poly-T oligo-attached magnetic beads. After fragmentation, the first strand cDNA was synthesized using random hexamer primers followed by the second strand cDNA synthesis. The library was ready after end repair, A-tailing, adapter ligation, size selection, amplification, and purification. The library was checked with Qubit and real-time PCR for quantification and bioanalyzer for size distribution detection. Strand specific library Messenger RNA was purified from total RNA using poly-T oligo-attached magnetic beads. After fragmentation, the first strand cDNA was synthesized using random hexamer primers. Then the second strand cDNA was synthesized using dUTP, instead of dTTP. The directional library was ready after end repair, A-tailing, adapter ligation, size selection, USER enzyme digestion, amplification, and purification. The library was checked with Qubit and real-time PCR for quantification and bioanalyzer for size distribution detection.

#### Clustering and sequencing

After library quality control, different libraries were pooled based on the effective concentration and targeted data amount, then subjected to Illumina sequencing on an Illumina NovaSeq X Plus sequencer.

#### Data quality control

Raw data (raw reads) of fastq format were first processed with fastq software. In this step, clean data (clean reads) were obtained by removing reads containing adapter or ploy-N as well as any reads associated with low quality raw data. At the same time, Q20, Q30 and GC content the clean data were calculated. All the downstream analyses were based on the clean data with high quality.

#### Reads mapping to the reference genome

Reference genome and gene model annotation files were downloaded from genome website. Use HISAT2 (2.2.1) to build the index of the reference genome, and use HISAT2 to align paired-end clean reads to 3 the reference genome (hg38 r_4.3.1).

#### Quantification of gene expression level

Gene-level read counts were quantified using featureCounts (v2.0.6), producing a matrix of read counts per gene per sample for downstream analysis performed in R (v4.x). Analysis was restricted to protein-coding genes. Genes with low expression were filtered by retaining genes with a total count greater than or equal to the number of samples.

#### Differential expression analysis

Differential expression analysis was performed using the DESeq2 R package (v1.42.0). DESeq2 provides statistical programs for determining differential expression in digital gene expression data using models based on negative binomial distribution. The resulting P-value is adjusted using the Benjamini-Hochberg methods to control the error discovery rate. The threshold of significant differential expression padj <= 0.05 and |log2(foldchange)| >= 1.

#### Transcription factor analysis of differentially expressed genes

Experimentally validated TF-target interactions of NFKB1 and LEF1 were obtained from the TFlink database and filtered to include only targets supported by small-scale evidence. Filtered target gene lists were intersected with the set of differentially expressed genes using gene symbols and genes present in both sets were defined as direct targets of the corresponding TF. For genes regulated by multiple TFs, fractional assignments were applied such that each gene contributed equally to each associated TF.

#### GO analysis of differentially expressed genes

Gene Ontology (GO) enrichment analyses were performed separately for upregulated and downregulated genes (defined as genes with adjusted p-values < 0.05 and log2 fold changes > 1 or < −1, respectively) using the clusterProfiler package (v4.8.1). Overrepresentation analysis was conducted for Gene Ontology Biological Process (GO:BP) terms using annotations from the org.Hs.eg.db database. The background gene universe was defined as all genes tested in the differential expression analysis after filtering, and gene identifiers were converted to Entrez Gene IDs prior to enrichment analysis. GO terms were considered significantly enriched at a Benjamini–Hochberg adjusted p-value < 0.05.

#### Hierarchical clustering

For visualization, gene expression values were transformed using the variance stabilizing transformation (vst) implemented in DESeq2 to account for mean-variance dependence in count data. The top 500 differentially expressed genes, ranked by adjusted P-value, were scaled by gene (row-wise z-score) and visualized using a hierarchical clustering heatmap generated with the pheatmap package.

### Proteomics

#### Sample preparation

Posterior tibial artery segments were obtained from LifeNet Health and flash frozen. 5mm segments of cellular and acellular vessels were flash frozen and prepared for liquid chromatography mass spectrometry by Arcproteomics using standard procedures.

#### Protein Digestion (Cell Pellet)

Samples were lysed with 50ul of 8M urea with protease and phosphatase inhibitors (ThermoFisher) and ice batch sonicated for 10 mins. EasyPep lysis buffer (50ul) was added and cup horn sonicated in chilled water. BCA was perform and 5 ug was used for digestion. Samples were reduced with DTT (5 mM) and alkylated with IAA (10mM) for 10mins at 80°C. The samples were then digested with 1ug of Lysyl endopeptidase (Wako) and 2ug trypsin (Thermofisher) overnight at room temperature. The peptide solutions were then acidified to 1:1 with 4% H_3_PO_4_ and cleaned with microelute MCX columns according to manufacturer’s protocol. The eluates were then dried to completeness using a SpeedVac (LabConco).

#### Liquid Chromatography and Mass spectrometry (Evosep ONE)

Each sample was resuspended in 100 ul of loading buffer (0.1% FA) and 20 ul was loaded onto Evotips and analyzed by liquid chromatography coupled to tandem mass spectrometry. Peptide eluents were separated on IonOptick’s column (15 cm × 75 μM internal diameter (ID) packed with 1.7um resin) by a Evosep One (Evosep). Buffer A was water with 0.1% (vol/vol) formic acid, and buffer B was 100% (vol/vol) acetonitrile in water with 0.1% (vol/vol) formic acid. Elution was performed using the preset 40SPD Whisper Zoom method. Peptides were monitored on a Orbitrap Astral mass spectrometer (ThermoFisher Scientific) fitted with a high-field asymmetric waveform ion mobility spectrometry (FAIMS Pro) ion mobility source (ThermoFisher Scientific). One compensation voltages (CV) of −35 was chosen for the FAIMS. Each cycle consisted of one full scan (MS1) was performed with an m/z range of 380-980 at 240,000 resolution, 500% AGC and 3 ms injection time. The higher energy collision-induced dissociation (HCD) DIA scans were collected with a 3 m/z isolation windows over the entire precursor range (380-980 m/z) with a time of 0.6 seconds and 3 ms injection time. Collision energy was set to 27% and scan range set to 150 - 2000 m/z.

#### Database Search

Spectronaut (version 19.9.250512.62635) was used to search all rawfiles in default library free mode using a human uniprot database (downloaded 012026 and supplemented with porcine collagen). All parameters were kept at default.

#### Data analysis

Protein intensity values were provided in log2-transformed form and imported into R for downstream analysis. Gene-level abundances were defined by aggregating the normalized log2 intensities using the median within each gene within each sample, and proteins detected in fewer than two samples per condition were excluded to reduce the influence of single-sample detections and improve robustness. Extracellular matrix (ECM) proteins were defined using the NABA_MATRISOME gene set (C2 curated gene sets) from the Molecular Signatures Database (MSigDB)^52,54^. Condition-specific protein universes were constructed and intersected with the matrisome gene set to identify ECM proteins present in each condition and to assess overlap between groups.

### Vasoconstriction

Vasoconstrictive responses were assessed using *ex vivo* cross-sectional imaging of engineered vessels. Following culture, vessels were manually sectioned into thin transverse slices (∼200 µm thickness). Individual slices were placed in Live Cell Imaging Solution (Gibco, A59688DJ) and oriented to visualize the cross-section of the lumen. The vessel slices were either imaged under basal conditions or 20μM Angiotensin II. Brightfield images were acquired immediately following stimulation and continuously for 180 seconds at a frame rate of 1 frame/second. Lumenal area was quantified by measuring thresholded pixel area within a manually defined inner-lumen ROI applied across 8-bit image stacks. A fixed global threshold was used, and white pixel area within the ROI was quantified over time using a custom ImageJ macro and normalized to a time zero baseline.

### FITC- Conjugated Dextran Permeability Experiments

Endothelial barrier function was evaluated using 25 μg/mL solutions of 20 kDa and 70 kDa FITC-conjugated dextran (info) in Cell Biologics Complete Endothelial Cell Medium Kit (Cell Biologics, H1168). Vessels were perfused at 20 μL/min for a total of 60 minutes on a heated live-cell imaging platform and visualized using a widefield fluorescent microscope (Zeiss Axio Observer 2). Images collected before perfusion (t_b_), at the start of perfusion (t_i_), and after 30 min (t) were analyzed using ImageJ (NIH, USA). Diffusional permeability was calculated using:

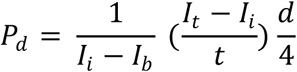

where P_d_ is the diffusional permeability index, I_b_ is the background intensity before perfusion, I_i_ is the average initial intensity, I_t_ is the average final intensity, and d is the artery diameter.

### Microscopy

Vasoconstriction and dextran permeability videos were acquired using a widefield light microscope (Zeiss AxioObserver 2, Hamamatsu ORCA-ER camera) with a Zeiss Plan-Achromat 4x/0.10NA objective.

Fluorescence confocal imaging was performed on a Zeiss Axio Observer 2 inverted microscope equipped with an LSM 880 confocal scan head, motorized z-stage, LSM T-PMT detectors, and 405, 488, 561, and 633 nm laser lines. Full vessel imaging was performed using tile scans and z-stack functionalities with a Zeiss EC Plan-Neofluor 10x/0.3 NA M27 objective. Higher magnification images were acquired with the Zeiss Plan-Apochromat 20x/0.8 NA M27 objective. All images were acquired using the Zeiss Zen Blue software and image processing was performed using FĲI/ImageJ 1.54f.

### Cytokine Immunoassays

Conditioned medium from perfusing vessels was collected at the indicated time points and frozen at −80°C until analysis. An ELISA to quantitate human IL-6 (DuoSets from R&D Systems) was performed per manufacturers’ instructions, with sample diluted to coincide within the range of standards. A custom human magnetic Luminex assay (R&D Systems, Minneapolis, MN) was performed per manufacturers’ instructions to measure cytokines present in the perfusate, including BAFF, CCL2, CCL20, CCL26, CXCL1, CXCL8, Pro-collagen 1a1, MMP-3, and MMP-9. Luminex assays were performed using the Luminex MAGPIX system with Luminex Exponent software (v4.2). Concentrations were calculated by a five-place logistic regression from standards within 80%-120% of the expected values.

### Statistics

Statistical analysis was performed using graphpad Prism with statistical tests as indicated. Processing and analysis of Luminex, transcriptomic, and proteomic data was performed with R (version 4.6.0) using packages including dplyr, tidyr, limma, DESeq2, clusterprofiler, msigdbr, ggplot2, and pheatmap.

### Data availability

RNAseq data sets are through GEO with accession number GSE335077

## ACKNOWLEDGEMENTS

We acknowledge AbbVie for funding and scientific support. We thank the Industrial Affiliates Program at the Institute of Material Science at the University of Connecticut for assistance with rheological and SEM material characterization. We thank the Yale Center for Cellular & Molecular Imaging for usage of their Zeiss LSM 880 AiryScan microscope. Finally, we thank Soumya Mitra, Annette Schwartz, and Prisca Honore for stimulating discussions and support.

## DISCLOSURES

SW is an employee of AbbVie. Financial support for this research was provided by AbbVie. AbbVie participated in the interpretation of data, review, and approval of the publication.

